# DNAJB8 oligomerization is mediated by an aromatic-rich motif that is dispensable for substrate activity

**DOI:** 10.1101/2023.03.06.531355

**Authors:** Bryan D. Ryder, Elizaveta Ustyantseva, David R. Boyer, Ayde Mendoza-Oliva, Mikołaj I. Kuska, Paweł M. Wydorski, Paulina Macierzynska, Nabil Morgan, Michael R. Sawaya, Marc I. Diamond, Harm H. Kampinga, Lukasz A. Joachimiak

## Abstract

J-domain protein (JDP) molecular chaperones have emerged as central players that maintain a healthy proteome. The diverse members of the JDP family function as monomers/dimers and a small subset assemble into micron-sized oligomers. The oligomeric JDP members have eluded structural characterization due to their low-complexity, intrinsically disordered middle domains. This in turn, obscures the biological significance of these larger oligomers in protein folding processes. Here, we identified a short, aromatic motif within DNAJB8, that drives self-assembly through ν-ν stacking and determined its X-ray structure. We show that mutations in the motif disrupt DNAJB8 oligomerization *in vitro* and in cells. DNAJB8 variants that are unable to assemble bind to misfolded tau seeds more specifically and retain capacity to reduce protein aggregation *in vitro* and in cells. We propose a new model for DNAJB8 function in which the sequences in the low-complexity domains play distinct roles in assembly and substrate activity.

**HIGHLIGHTS:** DNAJB8 oligomerization is mediated by a short phenylalanine-based motif in the S/T domain

Mutation of a single phenylalanine yields a monomeric form of DNAJB8

Monomeric DNABJ8 binds to an aggregation-prone substrate

Monomeric DNAJB8 retains substrate aggregation prevention activity

## INTRODUCTION

Cellular protein homeostasis is maintained by a complex network of molecular chaperones that regulate protein folding, refolding (Hartl 1996, Kampinga and Craig 2010, Ryder, Wydorski et al. 2022) and degradation (Fernandez-Fernandez, Gragera et al. 2017, Quintana-Gallardo, Martin-Benito et al. 2019). Disruption of this network increases the accumulation risk of proteins with folds that are prone to form proteotoxic β-sheet-rich amyloids, which are the hallmarks of many neurodegenerative diseases (Morawe, Hiebel et al. 2012, Hohn, Tramutola and Cascella 2020, Abrams, Arhar et al. 2021). For these amyloidogenic proteins, short sequence motifs contribute to the formation of aggregates (Sawaya, Sambashivan et al. 2007, Goldschmidt, Teng et al. 2010). The challenge for the cell is to discriminate protein conformations with these exposed motifs from non-pathogenic conformations that bury or sequester these motifs. Despite the critical role that chaperones play in preventing the formation of misfolded proteins, there are several lingering questions as to how they can discriminate between normal vs. disease conformations of substrates.

One of the primary chaperones involved in the cell proteostasis network is the highly conserved protein chaperone Hsp70, but one of the major weaknesses of this chaperone is its broad recognition of substrates (Kampinga and Craig 2010, Fernandez-Fernandez, Gragera et al. 2017, Rosenzweig, Nillegoda et al. 2019). This lack of specificity makes it a weak target in aggregation prevention. Given that there is a vast repertoire of possible unique sequence motifs that are exposed in amyloidogenic substrates, Hsp70 requires additional co-chaperones to facilitate specific recognition of motifs and substrates (Kampinga and Craig 2010, Ryder, Wydorski et al. 2022). This additional specificity is imparted by a family of J-domain proteins (JDPs) that classically coordinate substrate activities with Hsp70s (Hartl 1996, Glover and Lindquist 1998, Qiu, Shao et al. 2006, Kampinga and Craig 2010, Nillegoda, Kirstein et al. 2015). In the human proteome, JPDs are encoded by 47 known members and possess a diverse range of specialized domains for substrate recognition (Hartl 1996, Cheetham and Caplan 1998, Kampinga and Craig 2010, Faust, Abayev-Avraham et al. 2020, Piette, Alerasool et al. 2021, Zhang, Malinverni et al. 2023). Members of the JDP family bind to Hsp70 through a highly conserved J-domain (JD), which in turn triggers ATP hydrolysis within the nucleotide binding domain of Hsp70 and the closure of the substrate binding domain lid onto the protein substrate (Hartl 1996, Tsai and Douglas 1996, Kampinga and Craig 2010). The substrate is then refolded by Hsp70 and either restored to its native state or passed along for further processing by other chaperones such as Hsp90(Hartl 1996, Lackie, Maciejewski et al. 2017, Schopf, Biebl and Buchner 2017). The initial substrate recognition step by JDPs is crucial in the protein refolding network given the wide breadth of unique proteins in the cell. It is important that JDPs not only recognize unique protein sequences, but also identify irregular conformations that render proteins dysfunctional or potentially toxic. This is especially true for amyloidogenic proteins, where specific JDPs are able to recognize multiple distinct conformations or oligomeric states of a single protein substrate.

The microtubule-associated protein tau (herein; tau), which forms amyloids in a myriad of neurodegenerative diseases (Hong, Zhukareva et al. 1998, Clavaguera, Bolmont et al. 2009, Sanders, Kaufman et al. 2014, Goedert, Eisenberg and Crowther 2017, Ryder, Wydorski et al. 2022), is an example of one of these substrates. Tau is associated with several neurodegenerative diseases that are categorized as tauopathies (Sanders, Kaufman et al. 2014, Kaufman, Sanders et al. 2016). One of the most striking features of each tauopathy is that while all tauopathies are defined by the presence of tau aggregates in the brain, each tauopathy arises from several different structural polymorphs. Recently, cryo-EM has been used to solve the structures of several different tau fibril cores that are defined by arrangements of the tau repeat domains (RD)(Fitzpatrick, Falcon et al. 2017, Scheres, Zhang et al. 2020). As revealed by these structures, the variations in how the RDs are arranged lead to different sequence motifs becoming exposed or buried depending on the specific conformation. This means that a single protein recognition mechanism of tau is insufficient to recognize all possible polymorphs across different tauopathies. The wide diversity of the JDP family compensates for this complication, and provides motif-specific diversity in targeting aggregation-prone tau (Mok, Condello et al. 2018, Irwin, Faust et al. 2021).

There are a number of JDPs that have been shown to recognize different conformations of tau monomers or small oligomers (DNAJA1, DNAJA2, DNAJC5)(Burré, Sharma et al. 2010, Abisambra, Jinwal et al. 2012, Sharma, Burre et al. 2012, Mok, Condello et al. 2018, Irwin, Faust et al. 2021), as well as larger oligomers and fibrils (DNAJB1, DNAJB4)(Mok, Condello et al. 2018, Nachman, Wentink et al. 2020, Irwin, Faust et al. 2021). Recent work from our lab has shown that tau can adopt alternative aggregation-competent conformations (i.e. seeds) of monomer (Mirbaha, Chen et al. 2018, Hou, Chen et al. 2021) that are formed early in disease prior to the detection of oligomers and insoluble fibrils (Mirbaha, Chen et al. 2022). Derived from these studies, we showed that DNAJC7 preferentially interacts with natively folded conformations of tau over pathogenic mutants or monomeric seeds (Hou, Wydorski et al. 2021). However, the high sequence variability and diversity across different JDPs (Kampinga and Craig 2010) has made identifying JDPs that interact with different tau species difficult. Thus, it remains unknown how JDPs recognize the diversity of species adopted by a single protein.

A number of JDPs are poor candidates for use in classical structural biology methods because they often encode domains that are largely unstructured and dynamic (Kampinga and Craig 2010). Some JDPs assemble as dimers (DNAJA1/A2 and DNAJB1)(Irwin, Faust et al. 2021) while others remain monomeric. For a few, such as DNAJB6 and DNAJB8, the unstructured regions in some JDPs can lead to self-assembly and give rise to polydisperse oligomers (Hageman, Rujano et al. 2010, Gillis, Schipper-Krom et al. 2013, Månsson, Kakkar et al. 2014, McMahon, Bergink et al. 2021, Ryder, Matlahov et al. 2021). DNAJB6 and DNAJB8 evolutionarily belong to a distinct subgroup of the DNAJB1 subclass. Whilst encoding similar architectures that include an N-terminal JD and a flanking unstructured glycine/phenylalanine-rich (G/F) domain, the regions C-terminal of this G/F region are entirely different, with DNAJB6 and DNAJB8 containing a serine-/threonine-rich (S/T) domain and a CTD that can interact with the JD, thereby regulating recruitment of Hsp70(McMahon, Bergink et al. 2021, Ryder, Matlahov et al. 2021). Previous work has shown that in DNAJB8, phenylalanines in the G/F and S/T domains are required for its assembly into polydisperse oligomers (Hageman, Rujano et al. 2010, Ryder, Matlahov et al. 2021), which reach sizes larger than what is reported for DNAJB6 oligomers (Månsson, Kakkar et al. 2014). Additionally, we and others have shown that the folded JD and CTD can interact, thereby regulating recruitment of Hsp70(McMahon, Bergink et al. 2021, Ryder, Matlahov et al. 2021). It remains unclear what role oligomerization of DNAJB6 and DNAJB8 plays in their activity, but both were initially discovered as potent inhibitors of amyloid protein assembly (Gillis, Schipper-Krom et al. 2013, Månsson, Kakkar et al. 2014, Kakkar, Mansson et al. 2016, Aprile, Kallstig et al. 2017, Osterlund, Lundqvist et al. 2020, Arkan, Ljungberg et al. 2021), with DNAJB8 showing a slight advantage in cells (Gillis, Schipper-Krom et al. 2013).

Initial genetic studies showed that DNAJB6 is essential for prevention of poly-glutamine (poly-Q) aggregation of the Huntington protein (Htt)(Hageman, Rujano et al. 2010). Later studies showed that these oligomeric JDPs are active across a wide range of substrates including α-synuclein (Arkan, Ljungberg et al. 2021), Aβ(Osterlund, Lundqvist et al. 2020), TDP-43 and others (Aprile, Kallstig et al. 2017), (Udan-Johns, Bengoechea et al. 2014). Studies on DNAJB6 and DNAJB8 interactions with Htt uncovered that the S/T domain is essential for substrate activity and its deletion (Hageman, Rujano et al. 2010) or mutation (Kakkar, Mansson et al. 2016) interferes with Htt substrate activity. Studies to determine mechanism indicated that DNAJB6 may function to block nucleation of Htt aggregation consistent with chaperone activity at substoichiometric ratios (Kakkar, Mansson et al. 2016). Ultimately, the detailed mechanism of DNAJB6 and DNAJB8 substrate activity and the role of oligomerization in this process remain poorly understood.

Here, we used multi-disciplinary methods to understand the minimal driving forces that govern DNAJB8 self-assembly and how DNAJB8 oligomerization may influence substrate activity. We identified a ^147^AFSSFN^152^ motif within the S/T-rich domain that drives assembly. Mutations of F151 to serine in both the peptide and full-length (FL) DNAJB8 yields a loss of self-assembly. And yet, using a series of DNAJB8 mutants, we provide *in vitro* and biological evidence that self-assembly is not essential for the anti-amyloidogenesis functionality of DNAJB8. Our data provide mechanistic insight into the detailed mode of DNAJB8 assembly, allowing us to decouple JDP self-assembly and substrate activity.

## RESULTS

### Phenylalanine residues in the S/T-rich domain drive self-assembly

DNAJB8 contains four domains (Figure 1A): the N-terminal conserved JD (Figure 1A; red), the disordered G/F-rich (Figure 1A; blue) and S/T-rich (Figure 1A; cyan) domains, and a β-stranded auto-inhibitory CTD (Figure 1A; green). In a previous study, we mutated all 18 phenylalanines to serines in the DNAJB8 disordered G/F-rich and S/T-rich domains yielding a variant that is monomeric and unable to self-assemble (Ryder, Matlahov et al. 2021). We set out to understand how the different 18 phenylalanines in the G/F and S/T domains of DNAJB8 influence its self-assembly. We first considered the distribution and density of phenylalanines within the G/F-rich and S/T-rich domains. We find that the phenylalanines in the middle of the G/F-rich domain contains two dense clusters of phenylalanine residues, while the periphery of the G/F-rich and S/T-rich domains possess a less dense but more evenly distributed arrangement of phenylalanine residues (Figure 1A). Guided by this analysis, we divided the G/F and S/T domains into four segments (DNAJB8_S1_: F87S, F93S, F100S, F103S, F104S; DNAJB8_S2_: F110S, F112S, F114S, F119S, F134S; DNAJB8_S3_: F138S, F141S, F144S, F148S, F151S; and DNAJB8_S4_: F164S, F169S, F179S) to capture the phenylalanine clusters while maintaining similar numbers of phenylalanines in each segment (Figure 1A). We next tested the capacity of FL DNAJB8 S1-S4 mutants to self-assemble in cells and *in vitro* (Figure 1B).

**Figure 1.**
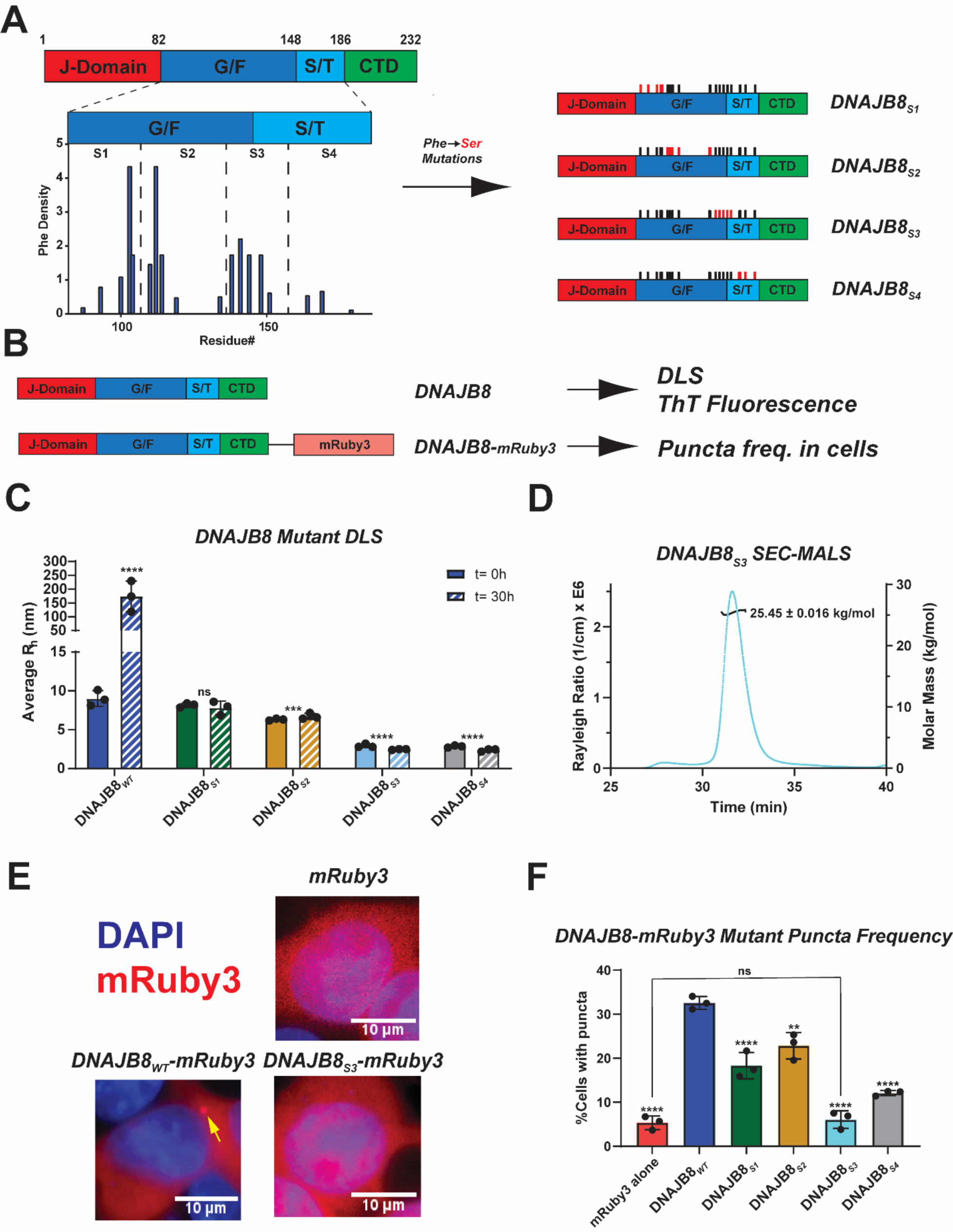
Phenylalanine residues in the S/T-rich domain drive self-assembly. (**A**) Domain map of DNAJB8 is colored according to domain annotation: JD (red), G/F-rich (blue), S/T-rich (cyan), and CTD (green). A histogram below the G/F-rich and S/T-rich domains shows the distribution of phenylalanine amino acids. Each position is given an arbitrary value to reflect local phenylalanine densities. Based on phenylalanie distributions and density characterizations, we arbitrarily divided the disordered domains into 4 segments (shown as S1, S2, S3, S4). We designed Phe→Ser substitution mutants (termed DNAJB8_S1_, DNAJB8_S2_, DNAJB8_S3_, and DNAJB8_S4_) for each segment with domain maps indicated which Phe residues (black bars) were mutated to Ser (red bars). (**B**) Domain maps for DNAJB8 used in cell experiments (DNAJB8-mRuby3) and *in vitro* experiments (DNAJB8). mRuby3 is colored pale red. (**C**) Average R_h_ measured by DLS of DNAJB8_WT_ (blue), DNAJB8_S1_ (green), DNAJB8_S2_ (gold), DNAJB8_S3_ (cyan), and DNAJB8_S4_ (grey) at time points t=0hrs (solid) and t=30hrs (dashed). DNAJB8_WT_ forms larger oligomers and assemblies after 30 minutes, while the other mutants remain at consistent sizes. While DNAJB8_S1_ and DNAJB8_S2_ primarily remain as larger oligomers such as trimers and tetramers, DNAJB8_S3_ and DNAJB8_S4_ remained at smaller sizes consistent with monomeric IDPs. Shown values were calculated with SOS < 20. For multiple comparisons to DNAJB8_WT_: *** p<0.001, and **** p<0.0001. (**D**) SEC-MALS chromatogram of DNAJB8_S3_. 99.3% of the sample mass eluted as a single peak with a calculated molar mass of 25.45 ±0.016 kg/mol. These results indicate that the protein is a stable monomer in solution. (**E**) Representative images from cells transfected with mRuby3 (top right), DNAJB8 (bottom left), and DNAJB8_S3_ (bottom right). DAPI is colored blue and mRuby3 signal is shown in red. DNAJB8 is expressed in the cytoplasm and forms cytoplasmic and juxtanuclear puncta. DNAJB8_S3_ shows few instances of puncta, but is able to be expressed in both the cytoplasm and nucleus. (**f**) Histogram of puncta frequency across cell lines expressing mRuby3 (red), DNAJB8_WT_-mRuby3 (blue), DNAJB8_S1_-mRuby3 (green), DNAJB8_S2_-mRuby3 (gold), DNAJB8_S3_-mRuby3 (cyan), and DNAJB8_S4_-mRuby3 (grey). Phenylalanine to serine mutations within the S/T-rich domain in DNAJB8_S3_ and DNAJB8_S4_ resulted in a decrease of puncta. All values were calculated from CellProfiler v.4.2.1. One-way ANOVA was used to report significance where F=75.25 and P-value<0.0001. For multiple comparisons to DNAJB8_WT_, ** p<0.01.

To determine if the serine mutations are changing the size and assembly of recombinant WT and mutant DNAJB8 *in vitro*, we used dynamic light scattering (DLS) to measure changes in the distribution of hydrodynamic radii (R_h_) over time. Due to the disordered G/F-rich and S/T-rich domains, DNAJB8 monomer and oligomer estimation was found to follow the particle size estimation model for intrinsically disordered proteins (Marsh and Forman-Kay 2010, Ryder, Matlahov et al. 2021) as opposed to the classical globular model. We observed that DNAJB8_WT_ demonstrated evidence of assembly into larger oligomers (>100nm) over 30 hours, with an initial average R_h_ of 7.9 ± 0.2 nm, indicating the presence of dimers and small oligomers (Figure 1C and Figure S1A; blue). This was comparable to the R_h_ distribution of DNAJB8_S1_ 8.8 ± 0.1 nm and DNAJB8_S2_ 7.3 ± 0.2 nm (Figure 1C and Figure S1A; green, gold). However, the average particle size of DNAJB8_S3_ and DNAJB8_S4_ were smaller at 2.4 ± 0.1 nm and 4.5 ± 0.6 nm, respectively (Figure 1C and Figure S1A; cyan, grey). These values are comparable to the size of monomeric DNAJB8 with all phenylalanines mutated to serine that we previously characterized (Ryder, Matlahov et al. 2021). Based on the initial sizes of each mutant, the similarity of DNAJB8_S1_ and DNAJB8_S2_ to DNAJB8_WT_ are due to their capacity to form dimers that lead to larger assemblies as we previously characterized (Ryder, Matlahov et al. 2021). By contrast, DNAJB8_S3_ and DNAJB8_S4_, remain small, consistent with sizes expected for a monomer (Marsh and Forman-Kay 2010) and consistent with a previously characterized DNAJB8 monomeric mutant (Ryder, Matlahov et al. 2021). Out of the four mutants, the DNAJB8_S3_ mutant size was further validated by size exclusion chromatography-multi angle light scattering (SEC-MALS). The SEC-MALS chromatogram showed that 99.3% of the total sample mass was a monomeric particle with a molar mass of 25.45 ± 0.016 kg/mol (Figure 1D), thus confirming that DNAJB8_S3_ is a stable monomer *in vitro*. These data show that the S/T-rich domain contains the specific phenylalanines that drive DNAJB8 oligomerization.

Recently, alternative classes of nonpolar motifs enriched in aromatic side chains have been implicated in protein self-assembly described as low complexity aromatic-rich kinked segments (LARKS)(Sawaya, Sambashivan et al. 2007, Hughes, Sawaya et al. 2018). These motifs stabilize β-sheet conformations that self-assemble through nonpolar side chain contacts and β-strand hydrogen bonds often detectible with Thioflavin T (ThT). DNAJB6, homolog to DNAJB8, is predicted to encode steric zipper motifs (Hughes, Sawaya et al. 2018). Based on this information, we hypothesized that a short, LARKS-like motif could be driving DNAJB8 oligomerization based on the evenly spaced arrangement of Phe residues in the S/T-rich domain, and the homology of these regions to proposed LARKS in DNAJB6b (Hughes, Sawaya et al. 2018) (Figure 1A). To consider this possibility, we used a ThT fluorescence aggregation assay to compare β-sheet-mediated assembly of our DNAJB8 mutants. DNAJB8_WT_ and DNAJB8_S1_ showed the highest ThT fluorescence signal at 181.3 ± 23.6 F.U. and 196.3 ± 15.2 F.U., respectively (Figure S1B; blue and green). DNAJB8_S2_ and DNAJB8_S4_ showed an approximate 2-fold decrease in fluorescence signal relative to WT with 104.0 ± 3.0 F.U. and 97.0 ± 9.9 F.U., respectively (Figure S1B; gold and grey). However, the monomeric DNAJB8_S3_ mutant did not yield any fluorescent signal, consistent with the inability to form β-sheet-rich oligomers (Figure S1B; cyan). These results support that mutations in DNAJB8_S3_ are indeed the most disruptive to DNAJB8 self-assembly, and further strengthen the hypothesis that a short sequence motif within the S/T domain may drive the self-assembly of DNAJB8 via β-sheet structures.

While our DLS, SEC-MALS, and ThT data have shown that DNAJB8 oligomerization can be disrupted by serine substitutions within the S/T-rich domain, the next question is whether or not this behavior translates into cells. One known marker of DNAJB8 assembly in cells is the presence of puncta (McMahon, Bergink et al. 2021, Ryder, Matlahov et al. 2021), which could be an indication of large oligomers or inclusions that interact with potential substrates. To test the effect of DNAJB8 serine segment mutants on puncta formation in cells, we expressed DNAJB8_WT_, DNAJB8_S1_, DNAJB8_S2_, DNAJB8_S3_, and DNAJB8_S4_ mutants as C-terminal fusions to mRuby3 in HEK293T cells. The cells were transfected, incubated for 48 hours at 37°C, and imaged on a high content microscope. After imaging, the cells were lysed and the lysate was run on native PAGE to observe relative abundance of higher molecular weight (oligomeric) DNAJB8-mRuby3 spices (Figure S1C). The images were analyzed to quantify the frequency, size and intensity of DNAJB8 puncta based on mRuby3 fluorescence (see methods). Quantification of fluorescence intensity showed even levels of protein mRuby3-based expression (Figure S1D) with no effects on viability (Figure S1E). Just as it was previously observed with a mClover3 fusion protein (Ryder, Matlahov et al. 2021), expression of DNAJB8_WT_-mRuby3 led to the formation of cytoplasmic and juxtanuclear puncta with an average diameter of 1.1 ± 0.3 μm with signal excluded from the nucleus (Figure 1E and Figure S1F). In general, F→S mutations within the disordered domains of DNAJB8 resulted in significant decreases in the number of puncta across cells (Figure 1F). DNAJB8_WT_ showed the highest number of puncta at 32.55 ± 1.46% (Figure 1F; blue), which is significantly higher than our negative control lines expressing only mRuby3 with 5.33 ± 1.56% (Figure 1F; red). When run on a native gel, DNAJB8_WT_ shows a wide range of different oligomeric species of low to high molecular weights (Figure S1C). While a similar number of puncta was observed for cells expressing DNAJB8_S1_ (18.3 ± 2.98%; Figure 1F; green) and DNAJB8_S2_ (22.83 ± 2.97%; Figure 1F; gold), there was a 5-fold and 3-fold decrease in the number of puncta in cells expressing DNAJB8_S3_ (6.06 ± 1.98%; Figure 1F; cyan) and DNAJB8_S4_ (12.09 ± 0.57; Figure 1F; grey), respectively. In parallel to the observed decreases in puncta, the cell lysates of DNAJB8_S3_ and DNAJB8_S4_ show stronger and more defined monomer bands on a native gel along with a loss of high molecular weight oligomers (Figure S1C). Out of all the F→S mutants, only DNAJB8_S3_ showed no statistically significant difference to the mRuby3 control (Figure 1F). Furthermore, while cells expressing DNAJB8 that encoded the S1, S2 and S4 mutations displayed the same cytoplasmic localization as DNAJB8_WT_-mRuby3 (Figure S1G-L), we observed that DNAJB8_S3_ was present in both the cytoplasm and the nucleus (Figure 1E and Figure S1K). Mutations of phenylalanine to serine in S3 segment reduce the number of DNAJB8 puncta, increase the abundance of monomers, and promote nuclear localization (Figure 1E-F and Figure S1C,K). These data show that as observed *in vitro*, phenylalanine to serine substitutions within the S/T-rich domain are changing how DNAJB8 assembles in the cell.

### A steric zipper motif drives local S/T-rich domain self-assembly

Based on the results of the cell and *in vitro* experiments, we identified the phenylalanine residues in DNAJB8 self-assembly are among 5 residues in segment 3: F138, F141, F144, F148, and F151. Out of these, F141, F144, F148, and F151 all correspond to homologous sequences in the S/T domain of DNAJB6b that are proposed to be potential LARKS (Hughes, Sawaya et al. 2018). Guided by the presence of detectible ThT signal in oligomeric DNAJB8 (Figure S1B), we designed a ThT aggregation-based screen of peptides across the disordered G/F and S/T domains (Figure 2A; Table S1). All peptides used in this experiment were non-overlapping 20 amino acids fragments and spanned from the beginning of the G/F domain to the end of the C-terminal domain. Each peptide was monitored for its capacity to self-assemble using a ThT fluorescence assay over 10 hours. Only one peptide, B8_5, within the screen showed evidence of self-assembly by ThT, which includes 3/5 phenylalanines found in segment 3 of DNAJB8 (Figure 2B). Negative stain TEM confirmed the presence of fibril-like assemblies for B8_5 (Figure 2C). Interestingly, sequence-based prediction of amyloid motifs (Goldschmidt, Teng et al. 2010) in the B8_5 fragment did not predict amyloidogenic properties (Figure S2A), but B8_5 does contain two homologous sequences to DNAJB6b that were predicted to form LARKS-mediated assemblies (Hughes, Sawaya et al. 2018). This suggests that this fragment may adopt assemblies distinct from previously described steric zippers (Sawaya, Sambashivan et al. 2007). Nevertheless, our experimental results indicate that a minimal motif found in segment 3 may be responsible for ordered self-assembly of DNAJB8.

**Figure 2.**
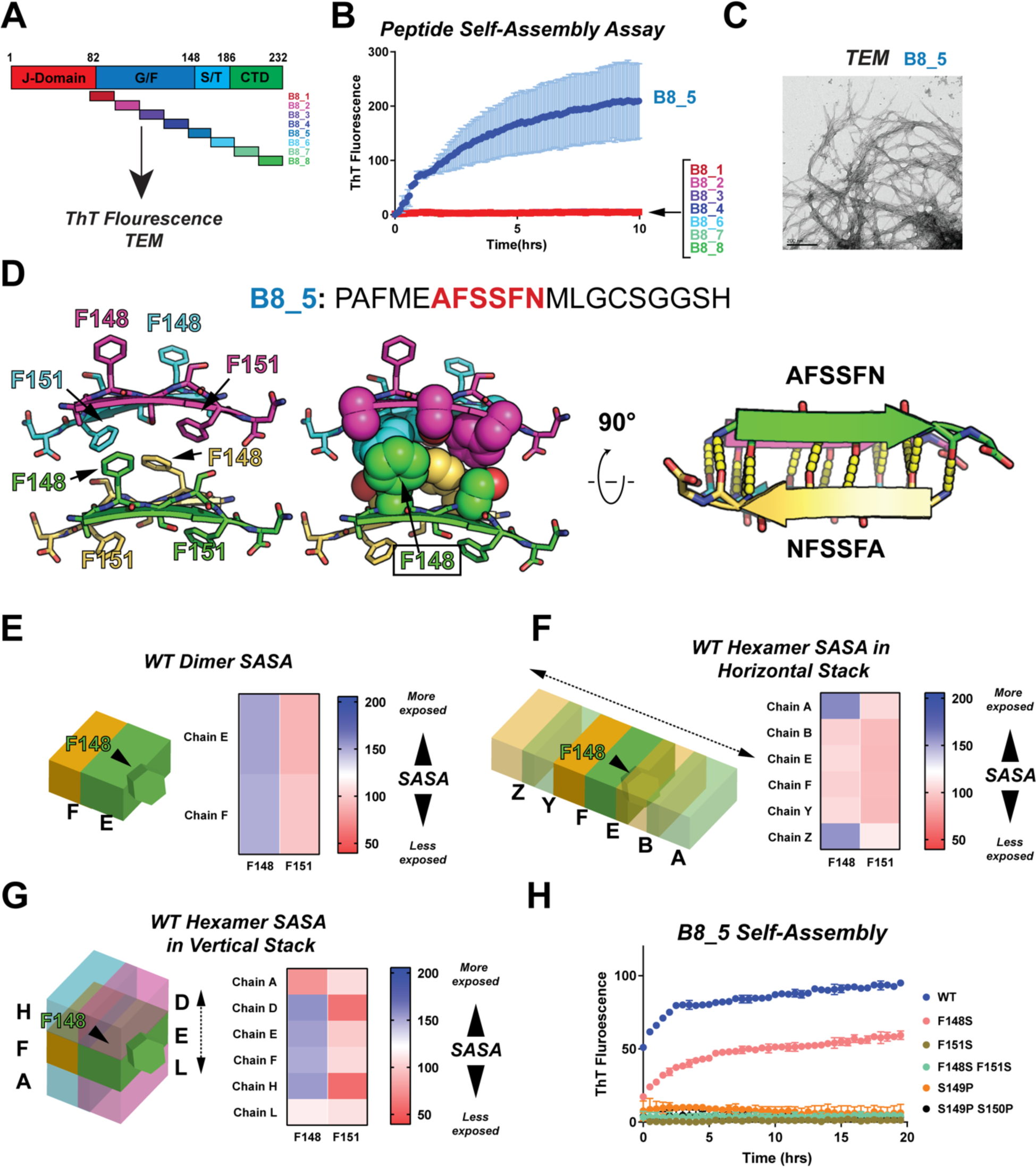
A steric zipper motif drives local S/T-rich domain self-assembly. (**A**) Domain map of DNAJB8 using the same color scheme as in Figure 1.1a. All peptides used in the peptide library are shown underneath their respective locations within the DNAJB8 sequence. (**B**) Thioflavin T time course experiment and TEM image of B8_5 fragment (blue). Out of all peptides used in the screen, only the B8_5 fragment showed evidence of assembly through increasing ThT fluorescence. Likewise, the B8_5 fragment was the only peptide to show visual evidence of large assemblies. Sale bar: 200nm. (**C**) TEM image of oligomeric B8_5 peptide assemblies after 10 hrs of incubation at room temperature. Scale bar: 200 nm. (**D**) 0.75Å microED structure of the ^147^AFSSFN^152^ class 6 steric zipper motif identified within the B8_5 peptide. Four peptides of ^147^AFSSFN^152^ are shown within the base subunit of the assembly (left). The structure is stabilized through hydrophobic packing of F148 and F151 (center), and anti-parallel β-stranded backbones (right). Crystallography statistics shown in Table S2. (**E**) SASA analysis of F148 and F151 in a peptide dimer based on the ^147^AFSSFN^152^ structure in panel d. A graphical representation shows the peptide arrangement used in the simulation. Higher SASA values correspond to greater solvent exposure (blue), while lower values correspond to less solvent exposure (red). (**F**) SASA analysis of F148 and F151 in a horizontally aligned hexameric assembly based on the ^147^AFSSFN^152^ structure in panel d. A graphical representation shows the peptide arrangement used in the simulation. Higher SASA values correspond to greater solvent exposure (blue), while lower values correspond to less solvent exposure (red). (**G**) SASA analysis of F148 and F151 in a vertically aligned hexameric assembly based on the ^147^AFSSFN^152^ structure in panel d. A graphical representation shows the peptide arrangement used in the simulation. Higher SASA values correspond to greater solvent exposure (blue), while lower values correspond to less solvent exposure (red). (**H**) Thioflavin T time course of B8_5 WT (blue) derived peptides where mutations are introduced to disrupt assembly based on Phe-Phe interactions: F148S (salmon), F151S (gold), F148S F151S (teal), or the β-strand backbone: S149P (orange), S149P S150P (black).

To gain structural insight into how DNAJB8 fragments may assemble, we generated a library of hexapeptides derived from the B8_5 fragment and screened them for the formation of crystals suitable for X-ray structure determination. Only one hexapeptide, ^147^AFSSFN^152^, yielded diffracting crystals. We determined a 0.75 Å resolution crystal structure of the ^147^AFSSFN^152^ assembly revealing a class 6 steric zipper motif (Sawaya, Sambashivan et al. 2007) (Figure 2D, Figure S2B, Table S2). The assembly is defined by two pairs of anti-parallel β-strands stabilized through hydrophobic interactions along opposite faces (face-to-back)(Sawaya, Sambashivan et al. 2007). The β-strand dimers show 2-fold symmetry within the assembly (Figure 2D and Figure S2C). The hydrophobic core of the assembly contains F151 on one dimer of β-strands surrounded on the edges by F148 from a second dimer. Within the core of the assembly, both F151 residues on a single dimer are oriented towards the center of the tetramer, but only one residue of F148 is oriented towards the center (Figure 2D). Interestingly, both F148 and F151 are positionally homologous with the predicted ^150^FGSGFS^155^ and ^157^FDTGFT^162^ LARKS in DNAJB6b, with a more direct alignment across both segments to ^153^GFSSFD^158^ (Hughes, Sawaya et al. 2018).

To distinguish which phenylalanine position is the most important in stabilizing the dimer, we minimized the structures in Rosetta (Mullapudi, Vaquer-Alicea et al. 2022) and calculated the solvent-accessible surface area (SASA) of each residue. The analysis was run in the context of the peptide dimer that was derived from the crystal symmetry (Figure 2E). We calculated the SASA for every residue across each peptide unit (labelled as “chains”). We first ran each chain as independent monomers and found that both F148 and F151 are relatively exposed (Figure S2E). In the context of the dimer, F148 remains exposed while F151 becomes more buried (Figure 2E; Figure S2F). When considering the orientation of the aromatic rings, we find that while the F151 pair interacts with each other across 2 chains to stabilize the structure, only one residue of F148 is oriented towards the center of the dimer, while the other is exposed to the side (Figure S2D, left). This leads to a lower SASA for F151, where both instances of F151 contribute to each other’s burial within the minimal unit, while F148 does not (Figure 2E; Figure S2F). This implies that F151 burial is an important interaction to stabilize intermolecular contacts within the dimer.

To analyze the importance of F148 and F151 in the context of oligomeric assemblies, we used the dimer as a base unit and expanded the assemblies in a horizontal and vertical direction. Upon expansion of the assembly in a horizontal direction, we find that both F148 and F151 are buried, where F148-F148 interactions are buried in the horizontal plane (Figure 2F and Figure S2D, right). Meanwhile, F151 burial is independent of this horizontal axis and does not require additional dimer-dimer interactions to stabilize itself (Figure 2F; Figure S2G). In the vertical expansion of the dimer assembly, F148 shows higher SASA than the horizontal assembly due to a greater exposure of the side chains to the solvent (Figure 2G; Figure S2H). However, partial burial is observed from the chains above and below due to the hydrophobic F151 face (Figure 2G; Figure S2D, center). Analysis of the different interfaces formed in the crystal lattice reveals differential burial modes of F148 and F151. The horizontal assembly is more dependent on the F148-F148 interactions, while the vertical assembly is more dependent on the formation of the F151 hydrophobic face. Therefore, we can conclude that F151 is crucially important for the initial dimer formation and subsequent oligomeric assemblies, while two possible modes of oligomeric assemblies could exist that are dependent on the aromatic ring orientation of F148.

### Burial of phenylalanine at position 151 dictates assembly of DNAJB8 fragments

Given the importance of phenylalanine burial in the different geometries of the steric zipper, we hypothesized that mutations to these positions would either weaken or disrupt the formation of this motif, leading to disruption of self-assembly. To test this, we introduced mutations to the B8_5 fragment to determine the role of aromatic side chains (F148S, F151S, F148S_F151S) and β-sheet conformation (S149P, S149P_S150P). As before, we used a ThT fluorescence aggregation assay to monitor self-assembly of mutant peptides compared to the WT B8_5 sequence (Figure 2H). The results showed that self-assembly of the B8_5 with the F148S mutation was reduced, and self-assembly was abolished with the F151S and F148S_F151S mutations (Figure 2H; salmon, gold, teal). Furthermore, all B8_5 peptides containing a single or double serine to proline mutation in B8_5 were not able to self-assemble suggesting that backbone contacts are important for oligomerization (Figure 2H; orange and black).

We next determined how phenylalanine to serine mutations may influence the total SASA of the dimer and the horizontal and vertical hexamers. The F148S, F151S, and F148S_F151S mutations were introduced into the ^147^AFSSFN^152^ structure using Rosetta and minimized as previously described (Mullapudi, Vaquer-Alicea et al. 2022). The ϕλSASA values were calculated by comparing the SASA between WT SASA and each mutant across the dimer, horizontal and vertical hexamer assemblies (Figure S2E). Compared to the WT sequence, the F148S mutation led to a loss of SASA in both the dimer and horizontal assemblies, while F151S resulted in a gain of SASA (Figure S2F, salmon and brown). Given the disengagement of F148 in the structure, it is expected that a mutation to this position would lead to a greater loss of SASA due to the loss of the side chain. However, the mutation to F151 does not result in such a loss since both F151 residues are more buried in the structure due to their greater role in stabilizing the packing of the zipper. The double mutant, F148S_F151S showed additive effects from both mutations, which resulted in a net loss within the dimer, but no change within the horizontal assembly (Figure S2F, green). However, when considering the effect of the mutations on the SASA of the vertical assembly, we observed an overall gain of SASA from both single mutants and the double mutant (Figure S2F, right). These results are more consistent with the hypothesis that both F148 and F151 participate in SASA burial to a certain degree within the larger peptide assembly if the directionality is vertical. However, one explanation that is consistent with our data is that while F148 plays a role in stabilizing larger oligomers, F151 is critical for the initial peptide dimer formation that precedes the initiation of self-assembly.

We further tested the homologous peptide sequences within DNAJB6b to see if the S/T domain sequences spontaneously self-assemble similar to the B8_5 fragment in DNAJB8. No similar mode of assembly was observed for DNAJB6b S/T domain fragments or CTD sequences that were previously implicated in DNAJB6b assembly (Karamanos, Tugarinov and Clore 2019, Karamanos, Tugarinov and Clore 2020) (Figure S2G). These data indicate that the observed steric zipper structure resulting from the ^147^AFSSFN^152^ motif is unique to DNAJB8 and not shared by the DNAJB6b homolog, with DNAJB6b self-assembly driven through other molecular forces. Overall, these data validate that the ^147^AFSSFN^152^ steric zipper motif controls B8_5 fragment assembly.

### Phenylalanine at position 151 is the primary driver of FL DNAJB8 self-assembly

We next asked whether mutations designed to disrupt the steric zipper motif formed by ^147^AFSSFN^152^ also inhibited assembly of FL DNAJB8 *in vitro* and in cells. To test this, recombinant FL DNAJB8 constructs containing F148S, F151S, or F148S_F151S (herein DNAJB8_F148S_, DNAJB8_F151S_, DNAJB8_F148S_F151S_, respectively) were purified and their assembly tested in ThT fluorescence and DLS assays *in vitro* (Figure 3A). As observed in the peptide experiments, DNAJB8_F148S_ yielded a similar fluorescence signal as DNAJB8_WT_, but DNAJB8_F151S_ and the double mutant DNAJB8_F148S_F151S_ showed a decrease in ThT signal with the double mutant approaching the same fluorescence amplitude as DNAJB8_S3_ (Figure S3A). The capacity of single and double mutants to self-assemble was next confirmed using DLS. The initial average R_h_ of DNAJB8_F148S_ was 13.9 ± 0.7 nm, which is comparable to WT (Figure 3C and Figure S3B; blue and lime). Furthermore, evidence of DNAJB8 oligomers was observed after 30 hours in both DNAJB8_F148S_ and DNAJB8_WT_ (Figure S3B; blue and lime). However, DNAJB8_F151S_ and DNAJB8_F148S_F151S_ showed an inability to self-assemble over time and remained monomeric at 2.5 ± 0.5 nm (Figure 3C and Figure S3B; purple) and 2.5 ± 0.05 nm (Figure 3C and Figure S3B; teal) respectively, comparable to DNAJB8_S3_ and the previously reported mutant with all phenylalanines mutated (Ryder, Matlahov et al. 2021). We further analyzed the DNAJB8_F148S_F151S_ mutant using SEC-MALS, which showed that 93% of the sample eluted as a monomeric protein with a molar mass of 25.8 ± 0.2 kg/mol (Figure 3C). Together, these data suggest that the F151S mutation disrupts β-strand-dependent self-assembly *in vitro*, with the inclusion of F148S yielding an additive effect. We next measured puncta formation of DNAJB8-mRuby3 encoding F148S, F151S, and F148S F151S mutations (herein DNAJB8_F148S_-mRuby3, DNAJB8_F151S_-mRuby3, and DNAJB8_F148S_F151S_-mRuby3, respectively) in HEK293T cells. After transfection, the cells were incubated for 48 hours at 37°C before image collection on a high content microscope and the images were analyzed as before. Cell lysates were collected and run on native PAGE as before to observe relative monomer and oligomer abundance (Figure S3C). The average mRuby3 fluorescence intensity was similar across all the variants tested and expression of the proteins did not affect cellular viability (Figure S3D-E). While DNAJB8_F148S_-mRuby3 remained primarily within the cytoplasm and similar to DNAJB8_WT_-mRuby3, we observed that DNAJB8_F151S_-mRuby3 and DNAJB8_F148S_F151S_-mRuby3 both showed a decrease in puncta and both mutants were now able to enter the nucleus, similar to DNAJB8_S3_-mRuby3 (Figure 3D). While decreases in the number of puncta were observed across all mutants, DNAJB8_F148S_-mRuby3 showed the smallest decrease with 19.72 ± 4.09% of cells with puncta (Figure 3E; lime), while DNAJB8_F151S_-mRuby3 and DNAJB8_F148S_F151S_-mRuby3 showed a decrease with 5.35 ± 0.27% and 6.05 ± 0.97% of cells with puncta, respectively (Figure 3E; purple and teal). Again, this decrease in puncta was observed in parallel to the loss of oligomers and increase in monomer signal for DNAJB8_F151S_-mRuby3 and DNAJB8_F148S_F151S_-mRuby3 (Figure S3C). Not only did the single mutant DNAJB8_F148S_ result in a decrease in puncta across the cell lines comparable to DNAJB8_S1_ and DNAJB8_S2_, but the single mutant DNAJB8_F151S_ and double mutant DNAJB8_F148S_F151S_ resulted in a decrease of puncta comparable to DNAJB8_S3_ (Figure 1F and Figure 3E) and the previously reported DNAJB8_F→S_ mutant (Ryder, Matlahov et al. 2021). Consistent with the peptide experiments, our results on FL DNAJB8 point mutants support the importance of F151 in DNAJB8 self-assembly, with evidence that additional phenylalanine to serine substitutions in tandem with F148 decrease the oligomeric propensity of DNAJB8, predominantly yielding monomers. The fact that this motif is found within the S/T-rich domain is further supported by previous data that indicated the importance of the S/T-rich domain in self-assembly (Hageman, Rujano et al. 2010, Kakkar, Mansson et al. 2016).

**Figure 3.**
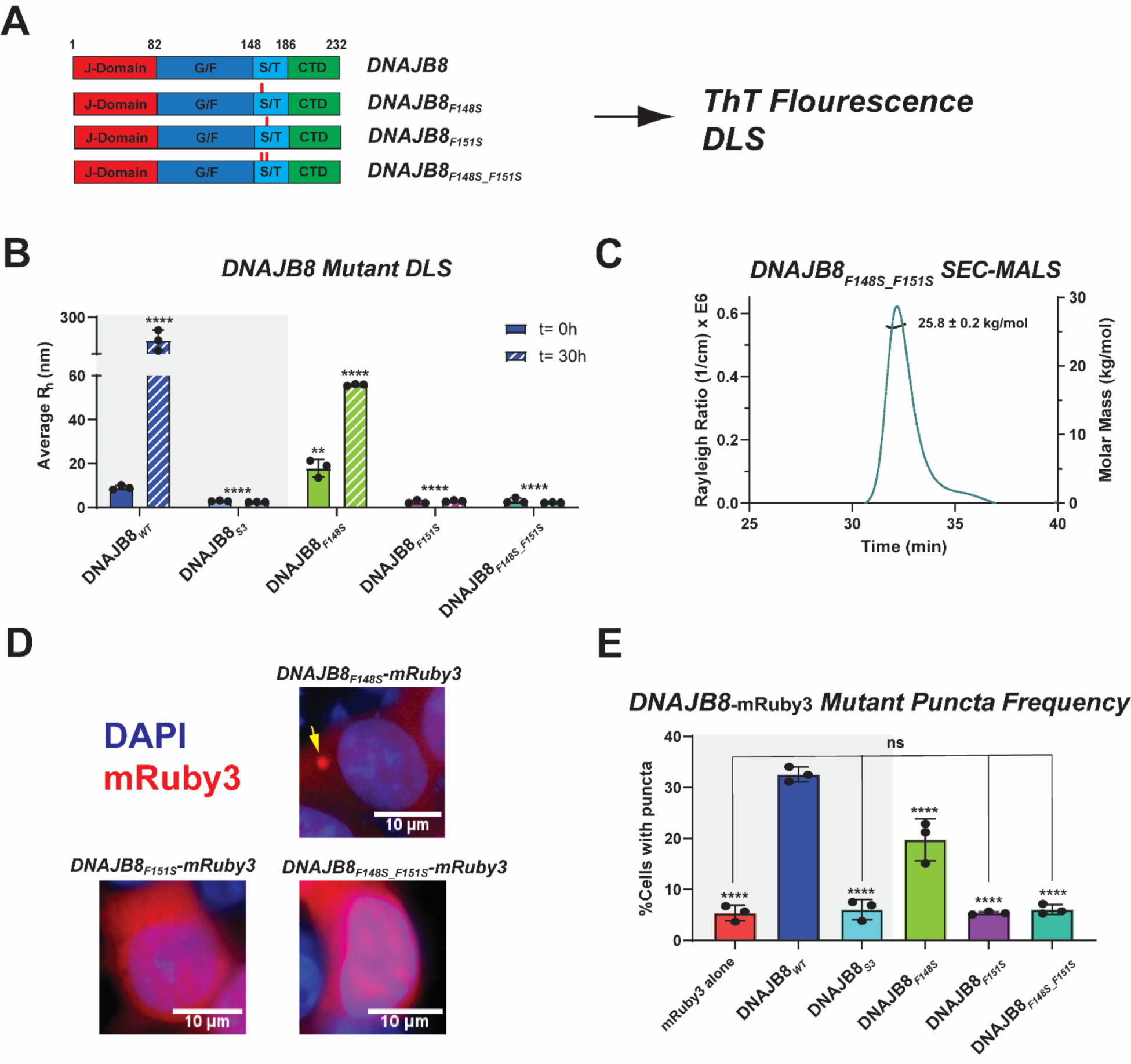
Phenylalanine at position 151 is the primary driver of FL DNAJB8 self-assembly. (**A**) Domain maps of DNAJB8_WT_, DNAJB8_F148S_, DNAJB8_F151S_, and DNAJB8_F148S_F151S_ using the same color scheme as in Figure 1a. Red hashes are shown to indicate the relative positions of the mutations. (**B**) Average R_h_ measured by DLS of DNAJB8_WT_ (blue), DNAJB8_S3_ (cyan), DNAJB8_F148S_ (lime), DNAJB8_F151S_ (purple), and DNAJB8_F148S_F151S_ (teal) at time points t=0hrs (solid) and t=30hrs (dashed). Data shown in the grey box was taken from Figure 1C for comparison. The average R_h_ of DNAJB8_F148S_ indicates the presence of larger oligomers in the sample at t=0 min, with some larger assemblies detected at t=30 min. Both DNAJB8_F151S_ and DNAJB8_F148S_F151S_ remained at a constant R_h_ consistent with monomeric IDPs. Shown values were calculated with SOS < 20. For multiple comparisons to DNAJB8_WT_: *** p<0.001, and **** p<0.0001. (**C**) SEC-MALS chromatogram of DNAJB8_F148S_F151S_. 93% of the sample mass eluted as a single peak with a calculated molar mass of 25.8 ±0.2 kg/mol. These results indicate that the protein is a stable monomer in solution. (**D**) Representative images of cell lines expressing DNAJB8_F148S_-mRuby3 (top right), DNAJB8_F151S_-mRuby3 (bottom left), and DNAJB8_F148S_F151S_-mRuby3 (bottom right). DAPI is shown in blue and mRuby3 signal from fusion proteins is shown in red. The single F148S mutation resulted in a cell population with less observed puncta than DNAJB8_WT_, with expression primarily in the cytoplasm and trace expression within the nucleus. However, the DNAJB8_F151S_ and DNAJB8_F148S_F151S_ mutants showed few instances of puncta and were expressed in both the nucleus and cytoplasm, sharing the same phenotype as DNAJB8 S3. (**E**) Histogram of puncta frequency across cell lines expressing mRuby3 (red), DNAJB8_WT_-mRuby3 (blue), DNAJB8_S3_-mRuby3 (cyan), DNAJB8_F148S_-mRuby3 (lime), DNAJB8_F151S_-mRuby3 (purple) and DNAJB8_F148S_F151S_-mRuby3 (teal). Data for mRuby3, DNAJB8 _WT_-mRuby3, and DNAJB8_S3_-mRuby3 (gray box) were previously shown in Figure 1F. Single and double phenylalanine to serine mutations at positions F148 and F151 decreased the abundance of puncta within each cell line, with the double mutant F148S/F151S reaching comparable levels to DNAJB8_S3_. This shows a relative decrease in self-assembly, with complete loss observed in DNAJB8_F151S_ and DNAJB8_F148S_F151S_. All values were calculated from CellProfiler v.4.2.1. One-way ANOVA was used to report significance where F=87.85 and P-value<0.0001. Significance relative to DNAJB8_WT_ reported as **** p<0.0001. Bar shown to indicate no significance between the mRuby3 negative control and DNAJB8_S3_, DNAJB8_F151S_, and DNAJB8_F148S_F151S_.

### DNAJB8 preferentially binds a misfolded tau seed

Previous work our lab characterized a minimal unit of tau called a “tau seed” (M_s_) that is capable of inducing recombinant tau aggregation (Mirbaha, Chen et al. 2018, Hou, Chen et al. 2021, Mirbaha, Chen et al. 2022). Comparison of the of the seeding monomer (M_s_) and normal inert tau (M_i_) ensembles uncovered differences in local interactions but also folding of the N-terminal acidic tail (Hou, Chen et al. 2021, Mirbaha, Chen et al. 2022). We have also previously developed assays to measure chaperone activity against both M_i_ and M_s_ forms of tau to show differences in DNAJC7 binding and activity (Hou, Wydorski et al. 2021). While oligomeric JDPs, such as DNAJB6 and DNAJB8, have not previously been reported to interact with tau, other class B JDPs are known to interact with tau fibrils (Mok, Condello et al. 2018, Faust, Abayev-Avraham et al. 2020, Irwin, Faust et al. 2021). Here, we tested directly whether DNAJB8_WT_ or DNAJB8_S3_ can discriminate between M_i_ and M_s_ to influence tau aggregation.

One of the most challenging complications in measuring DNAJB8_WT_ substrate binding affinity is the difficulty to parse out whether monomers or oligomers are binding to substrates. However, our DNAJB8_S3_ mutant remains monomeric, which eliminates this problem. Using this advantage, we first leveraged our engineered monomeric DNAJB8_S3_ to measure binding to M_i_ and M_s_ using microscale thermophoresis (MST) (Figure 4A). Titration of unlabeled M_i_ against Cy5 labeled DNAJB8_S3_ yielded a K_d_ > 400 μM indicating that the binding is very weak (Figure 4B, black and Figure S4A). Similarly, we also measured binding between M_s_ and DNAJB8_S3_ yielding a substantially tighter K_d_ of 24 μM (Figure 4B, red and Figure S4B). This change in affinity may be caused by differences in M_i_ and M_s_ conformations as was previously shown (Mirbaha, Chen et al. 2018, Chen, Drombosky et al. 2019, Hou, Chen et al. 2021), but most importantly, DNAJB8 is showing preferential binding to the aggregation-prone conformation of tau.

**Figure 4.**
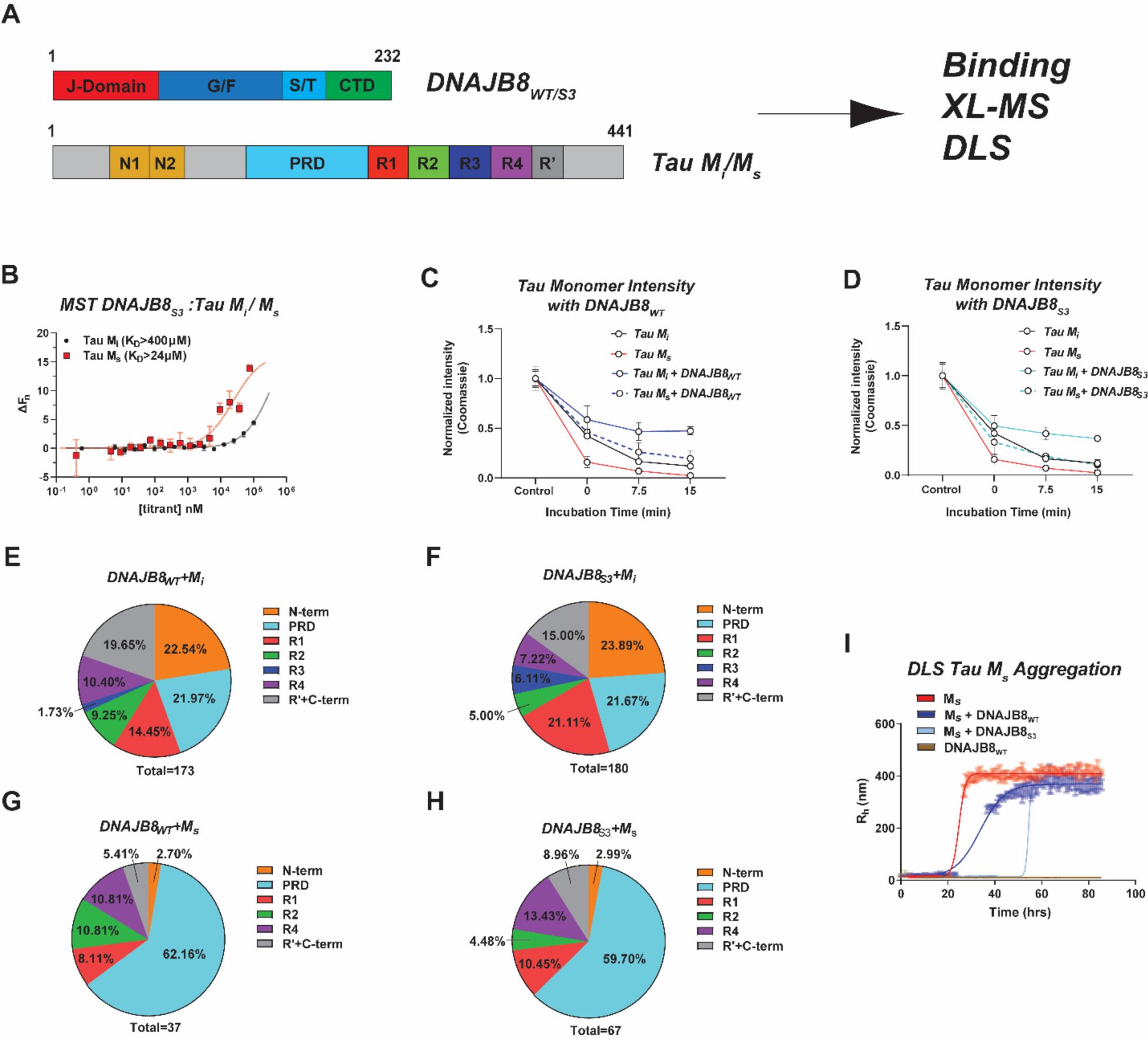
DNAJB8 preferentially binds a misfolded tau seed. (**A**) Domain maps of DNAJB8_WT_/DNAJB8_S3_, and full-length tau M_i_/M_s_ used in *in vitro* activity experiments. Domain color scheme for DNAJB8 is the same as shown in Fig 1a. Tau domains are shown from N to C termini as N-terminal domains (N1 and N2; orange), proline rich domain (PRD; cyan), and all repeat domains (R1; red, R2; green, R3; blue, R4; purple, R’; dark grey). (**B**) MST binding curves for S3 B8 binding to M_i_ tau (black, circles) or M_s_ tau (red, squares). Fn values calculated from thermophoretic time traces, as a function of titrated protein. Baseline for unbound thermophoresis has been subtracted for the clarity of representation. Error bars represent the range of three replicates. K_d_ difference between both tau species is ∼17-fold. (**C**) Relative coomassie signal intensity of the tau monomer band on a SDS-PAGE gel where DMTMM was added and quenched immediately (0 min), after 7.5 min, or after 15 min and either in samples of 1μM tau M_i_ (black), 1μM tau M_s_ (red), 1μM tau M_i_ with 20μM DNAJB8_WT_ (blue; solid), or 1μM tau M_s_ with 20μM DNAJB8_WT_ (blue;dashed). (**D**) Relative coomassie signal intensity of the tau monomer band on a SDS-PAGE gel where DMTMM was added and quenched immediately (0 min), after 7.5 min, or after 15 min and either in samples of 1μM tau M_i_ (black), 1μM tau M_s_ (red), 1μM tau M_i_ with 20μM DNAJB8_S3_ (cyan;solid), or 1μM tau M_s_ with 20μM DNAJB8_S3_ (cyan;dashed). (**E**) Contribution of different tau domains in the DNAJB8_WT_:tau M_i_ XL-MS low-resolution (id score >10) results. Out of a total 173 pairs, the majority of cross-links are evenly distributed across the N-terminal domain, PRD, R1, and C-terminal domains. This is indicative of a mix of different interactions across multiple states of tau M_i_. (**F**) Contribution of different tau domains in the DNAJB8_S3_:tau M_i_ XL-MS low-resolution (id score >10) results. Out of a total 180 pairs, the majority of cross-links are evenly distributed across the N-terminal domain, PRD, R1, and C-terminal domains. These results closely mirror the tau domain contributions when crosslinked to DNAJB8_WT_, showing a similar mode of interaction. (**G**) Contribution of different tau domains in the DNAJB8_WT_:tau M_s_ XL-MS low-resolution (id score >10) results. Out of a total of 37 crosslinks, a majority of 62.12% localize to the PRD, with contributions from all other domains being minimal. The decrease in unique contacts combined with the increased frequency around a specific domain indicates that the DNAJB8_WT_:tau M_s_ interaction is more defined than the DNAJB8_WT_: tau M_i_ interaction. (**H**) Contribution of different tau domains in the DNAJB8_S3_:tau M_s_ XL-MS low-resolution (id score >10) results. Out of a total of 37 crosslinks, a majority of 59.7% localize to the PRD, with contributions from all other domains being minimal. (**I**) Dynamic light scattering time course experiment reporting average R_h_ as time points for DNAJB8_WT_ (brown), tau M_s_ (red), DNAJB8 _WT_ and tau M_s_ (blue), and DNAJB8_S3_ and tau M_s_ (cyan). After 20 hrs, tau M_s_ is able to rapidly aggregate *in vitro* at the specified concentration of 0.1μM. Meanwhile, at 10μM concentrations, DNAJB8_WT_ is unable to self-assemble within the time course of the experiment. Each DNAJB8 mutant was mixed with tau M_s_ at a 100:1 ratio. At these conditions, DNAJB8_WT_ is able to slow the rate of tau aggregation by ∼20 hrs. However, the presence of DNAJB8_S3_ with tau M_s_ delays the onset of tau M_s_ assembly by ∼30 hrs.

Based on these data, we were interested in understanding whether the WT and mutant JDP may influence assembly of different forms of tau. We developed an assay to monitor crosslinking-based assembly of FITC-labeled M_i_ (FITC-M_i_) tau and quantify whether DNAJB8 can influence this process. By treating reactions with a crosslinker, depletion of monomer and formation of covalent adducts can be visualized on an SDS-PAGE gel. Based on prior crosslinking experiments on tau (Chen, Drombosky et al. 2019, Hou, Chen et al. 2021) and DNAJB8(Ryder, Matlahov et al. 2021), we optimized the reactions using 4-(4,6-dimethoxy-1,3,5-triazin-2-yl)-4-methyl-morpholinium chloride (DMTMM) which traps contacts between primary amines and carboxylates. 1μM FITC-M_i_, 20μM DNAJB8_WT_ and FITC-M_i_:DNAJB8_WT_ were treated with crosslinker for 0, 7.5, and 15 minutes, quenched and visualized on an SDS-PAGE gel. DNAJB8_WT_ shows a slow reduction of monomer as crosslinker incubation time increases, consistent with its capacity to self-assemble (Figure S4C-D). Meanwhile, M_i_ in the presence of DMTMM shows evidence of rapid assembly, with monomers being depleted rapidly into larger oligomers (Figure S4C-D). Quantification of the FITC-M_i_ band intensities reveals a near 10-fold depletion of tau (Figure 4C; black). By contrast, addition of DNAJB8_WT_ to M_i_ slows down the depletion of M_i_ by 5-fold (Figure 4C and Figure S4C-D). We also performed a parallel experiment with FITC-M_i_:DNAJB8_S3_. Consistent with DNAJB8_S3_ being monomeric, reactions containing only the mutant JDP, show little assembly (Figure S4E). Despite the inability to assemble, co-incubation of FITC-M_i_ with DNAJB8_S3_ yielded a 6-fold reduction of M_i_ depletion (Figure 4D). The close similarities in tau monomer retention between DNAJB8_WT_ and DNAJB8_S3_ show that both JDPs are active in our M_i_ tau assembly assay despite weak affinities.

To gain more resolution into the surfaces involved in the interactions between the JDP and tau we employed crosslinking mass spectrometry. While other structural approaches are not suitable for low affinity interactions, crosslinking mass spectrometry (XL-MS) can uncover binding surfaces for even weak interactions (Leitner, Faini et al. 2016, Ryder, Matlahov et al. 2021). In addition to probing interactions between DNAJB8_WT_ or DNAJB8_S3_ and M_i_, we also probed interactions between the two chaperones and tau M_s_. Triplicate samples were analyzed by mass spectrometry and processed using the Xquest pipeline (Walzthoeni, Joachimiak et al. 2015, Leitner, Faini et al. 2016). In the DNAJB8_WT_:M_i_ and DNAJB8_S3_:M_i_ containing data, the JD and CTD interact with different tau domains indiscriminately, suggesting weak non-specific interactions (Figure 4E-F; Figure S4G-H). By contrast, in the DNAJB8_WT_:M_s_ and DNAJB8_S3_:M_s_ datasets the patterns of interaction are different with both chaperones preferentially crosslinking to the PRD domains (Figure 4G-H; Figure S4I-J). Calculating the frequency of the contacts as a function of the entire dataset shows that the percentage of contacts to the PRD in the M_i_ experiments is 20% and increases to 60% in the M_s_ datasets (Figure 4E-H). Our XL-MS data support that M_s_ is folded differently than M_i_ consistent with our prior work (Chen, Drombosky et al. 2019) and that this change in conformation leads to more specific contacts between the JDPs (both WT and S3) to M_s_.

To test whether we can observe aggregation suppression of tau in the presence of DNAJB8_WT_, we used a thioflavin T fluorescence-based aggregation assay where a recombinantly derived minimal construct of tau containing only the repeat domains (tauRD) is seeded using tau M_s_. Our results show that at substoichiometric concentrations, tau M_s_ induces rapid tauRD aggregation (Figure S4K; orange, t_1/2max_= 6.3 ± 0.2 hours). Addition of DNAJB8_WT_ slows down M_s_-seeded tauRD aggregation nearly 10-fold (Figure S4K; black, t_1/2max_= 61.6 ± 2.9 hours). While these results show that the presence of DNAJB8_WT_ does delay tau aggregation, it is difficult to comment on how DNAJB8:M_s_ interactions specifically effect the nucleation stages of an aggregation experiment. Therefore, to test whether the chaperones can influence M_s_ self-assembly directly, we designed a DLS-based assay using low concentrations of M_s_ monomers (0.1μM) that are invisible to detect by scattering but once they form assemblies they scatter and can be detected directly. This experiment also removes M_i_ from the system, and unlike a ThT assay, directly reports on the isolated M_s_ conformation assembly at a concentration where ThT signal cannot be observed. Importantly, parallel reactions with 10μM WT and mutant JDPs remain flat in the assay allowing us to quantify their effect on M_s_ assembly by comparing the t_1/2max_ of the reactions (Figure 4I). M_s_ alone assembled with a t_1/2max_ of 24.7 ± 0.2 hours (Figure 4I, red), addition of DNAJB8_WT_ delays the increase in M_s_ assembly with a t_1/2max_ of 33.8 ± 0.3 hours (Figure 4I, blue). By contrast, reactions that included DNAJB8_S3_ delayed the M_s_ assembly even further yielding a t_1/2max_ of 54.3 ± 7.5 hours (Figure 4I, light blue). The addition of other mutants DNAJB8_S1_, DNAJB8_S2_, and DNAJB8_S4_ delayed the increase of M_s_ assembly similarly to DNAJB8_WT_ (Figure S4L). Since DNAJB8_S3_ is a monomer at these concentrations, these results show that self-assembly is not a pre-requisite for DNAJB8 to act as a holdase.

### Engineered monomeric DNAJB8 mutants retain anti-aggregation substrate activity in cells

Our data show that DNAJB8 is able to influence aggregation of tau *in vitro* despite weak binding affinities. We next tested whether our engineered DNAJB8 mutants are active against substrate aggregation in cells using tau and Htt as model substrates. We first used a cellular model of tau aggregation that co-expresses P301S tauRD-mCerulean3 and P301S tauRD-mClover3 in HEK293T cells (Figure 5A). Transduction of tau aggregates into these cells induces aggregation of the intracellular tau which can be quantified using FRET_Clover/Cerulean_(Holmes, Furman et al. 2014, Sanders, Kaufman et al. 2014, Hitt, Vaquer-Alicea et al. 2021). To test the effect of WT and mutant DNAJB8 on tau aggregation in this system, we co-transfected the tau biosensor cells with WT and mutant DNAJB8 C-terminally fused to mRuby3, followed by seeding the cells with pre-formed tau fibrils using lipofectamine (Figure 5A). We next interpreted changes in tau aggregation as measured by FRET_Clover/Cerulean_ but also changes in DNAJB8 oligomerization state as described above (Figure 1F and 3E). Parallel experiments were set up for imaging analysis and flow cytometry. After 48 hours following tau fibril treatment, one plate of cells was imaged to analyze tau and DNAJB8 inclusions, while a second plate was analyzed by flow cytometry to quantify tau FRET_Clover/Cerulean_ in each condition. Using DRAQ5 for nuclear staining, we were able to produce images showing DRAQ5, tau-mClover3, and DNAJB8_WT_-mRuby3/DNAJB8_S3_-mRuby3 for seeded and unseeded conditions (Figure S5A). In unseeded cells, the tau expression is diffuse throughout the cells (Figure S5A). Treatment of cells with 20nM fibrils leads to the formation of mClover3 positive tau puncta (Figure S5A). Consistent with our earlier experiments in HEK293T cells expressing only the chaperone, cytoplasmic DNAJB8 puncta were observed in cells expressing DNAJB8_WT_-mRuby3 (Figure S5A; yellow arrows), while DNAJB8_S3_-mRuby3 remained diffuse in the tau biosensor cell lines. In cells that have both DNAJB8_WT_-mRuby3 and tauRD-mClover3 puncta, the two inclusions do not colocalize (Figure S5A).

**Figure 5.**
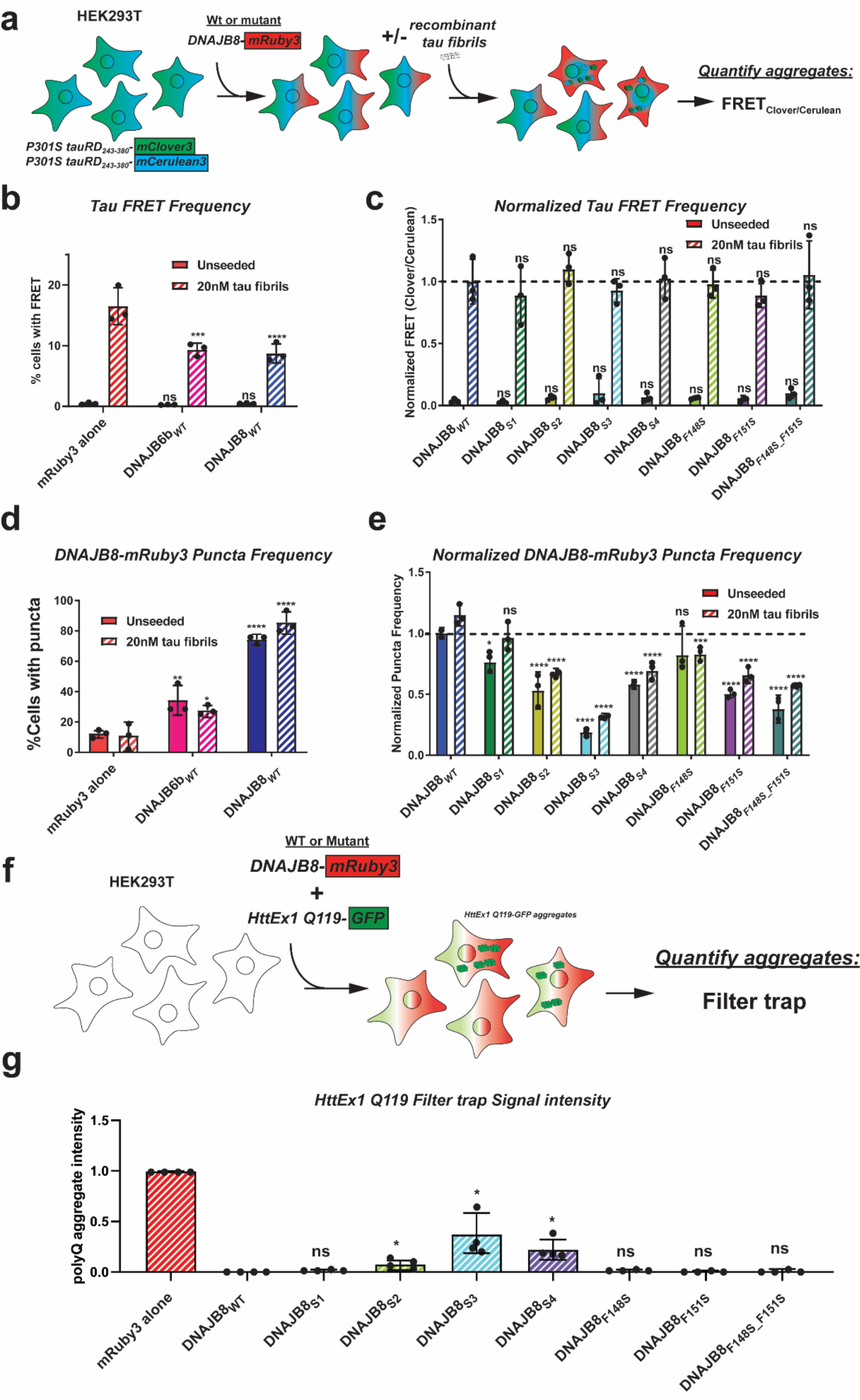
Engineered monomeric DNAJB8 mutants retain anti-aggregation substrate activity in cells. (**A**) Schematic of experiments involving stable HEK293T P301S tau biosensor lines. The tau biosensor cells are a stable line endogenously co-expressing P301S tauRD-mClover3 and P301S tauRD-mCerulean3. Upon transient transfection with sonicated tau fibrils, aggregation can be quantified through FRET_Clover/Cerulean_. Before adding sonicated fibrils, DNAJB8-mRuby3 constructs are first transfected into the stable cell lines for expression 24 or 72 hours before seeding with fibrils. Changes to tau aggregation are then quantified by FRET_Clover/Cerulean_. (**B**) Tau biosensor lines where transfected with mRuby3 (red), DNAJB6b_WT_-mRuby3 (pink), and DNAJB8_WT_-mRuby3 (blue) and expressed for 24 hours before either treatment with lipofectamine (solid bars) or seeded with 20nM sonicated tau fibrils (dashed lines). These lines were prepared as biological triplicate populations. Aggregation was quantified as a percentage of the cell line with positive FRET_Clover/Cerulean_ signal. Relative to the mRuby3 control, both cell lines expressing DNAJB6b_WT_ or DNAJB8_WT_ showed a 2-fold decrease in FRET_Clover/Cerulean_ positivity within the cell populations. (**C**) Tau biosensor lines where transfected with DNAJB8_WT_-mRuby3 (blue), DNAJB8_S1_-mRuby3 (green), DNAJB8_S2_-mRuby3 (yellow), DNAJB8_S3_-mRuby3 (cyan), DNAJB8_S4_-mRuby3 (grey), DNAJB8_F148S_-mRuby3 (lime), DNAJB8_F151S_-mRuby3 (purple), and DNAJB8_F148S_F151S_-mRuby3 (teal) and expressed for 24 hours before either treatment with lipofectamine (solid bars) or seeded with 20nM sonicated tau fibrils (dashed lines). % FRET_Clover/Cerulean_ was normalized to DNAJB8_WT_ as shown in panel 5b to directly compare DNAJB8 mutants to DNAJB8_WT_ activity (dashed line). Two-way ANOVA was used to calculate statistics across unseeded and 20nM tau fibrils lines, where no significant changes could be reported across the different mutants for either condition. (**D**) Percentage of cells containing mRuby3 puncta across tau biosensor cells transfected with mRuby3 (red), DNAJB6b_WT_-mRuby3 (pink), and DNAJB8_WT_-mRuby3 (blue) and expressed for 24 hours before either treatment with lipofectamine (solid bars) or seeded with 20nM sonicated tau fibrils (dashed lines). A 5-fold increase of puncta is observed for DNAJB8_WT_-mRuby3 compared to the mRuby3 control under both unseeded and seeded conditions, indicating that this effect is driven by DNAJB8_WT_. Statistics were calculated using two-way ANOVA, where * p<0.1, ** p<0.01, **** p<0.0001. Values taken from image analysis with CellProfiler v.4.2.1. (**E**) Normalized number of puncta within tau biosensor cell populations transfected with DNAJB8_WT_-mRuby3 (blue), DNAJB8_S1_-mRuby3 (green), DNAJB8_S2_-mRuby3 (yellow), DNAJB8_S3_-mRuby3 (cyan), DNAJB8_S4_-mRuby3 (grey), DNAJB8_F148S_-mRuby3 (lime), DNAJB8_F151S_-mRuby3 (purple), and DNAJB8_F148S_F151S_-mRuby3 (teal) and expressed for 24 hours before either treatment with lipofectamine (solid bars) or seeded with 20nM sonicated tau fibrils (dashed lines). The data were normalized to DNAJB8_WT_-mRuby3 as shown in panel 5d to directly compare each mutant DNAJB8 construct to DNAJB8_WT_-mRuby3 (dashed line). Similar to HEK293T cell experiments in Figure 1d and Figure 3e, monomeric DNAJB8 mutants DNAJB8_S3_, DNAJB8_F151S_, and DNAJB8_F148S_F151S_ show a significant decrease in puncta frequency that is dependent on the DNAJB8 mutations, but no changes observed due to the effect of tau aggregation. Statistics were calculated using two-way ANOVA, where * p<0.1, ** p<0.01, *** p<0.001, **** p<0.0001. Values taken from image analysis with CellProfiler v.4.2.1. (**F**) Schematic of experiments involving co-expression of HttEx1 Q119-GFP with WT and mutant forms of DNAJB8-mRuby3 in HEK293T cells. HttEx1 Q119-GFP construct were co-transfected with WT and mutant DNAJB8-mRuby3 into HEK293T cells. The cells were lysed and HttEx1 Q119-GFP aggregates were quantified via a filter trap assay. (**G**) FTA results showing HttEx1 Q119 aggregation for cells expressing mRuby3 (red), DNAJB8_WT_-mRuby3 (blue), DNAJB8_S1_-mRuby3 (green), DNAJB8_S2_-mRuby3 (yellow), DNAJB8_S3_-mRuby3 (cyan), DNAJB8_S4_-mRuby3 (grey), DNAJB8_F148S_-mRuby3 (lime), DNAJB8_F151S_-mRuby3 (purple), and DNAJB8_F148S_F151S_-mRuby3 (teal). All values were normalized to the mRuby3 negative control. A small loss of DNAJB8 activity was reported for DNAJB8_S2_-mRuby3, DNAJB8_S3_-mRuby3, and DNAJB8_S4_-mRuby3 expressing cells. Statistics were calculated using Welch’s T test for each pair relative to DNAJB8_WT_, where * p<0.1.

To evaluate the effect of DNAJB8 variants on tau seeding, we quantified the FRET_Clover/Cerulean_ signal in mRuby3 positive cells (Figure S5B). Cell lines not treated with tau fibrils had no detectible FRET_Clover/Cerulean_ (Figure 5B). For cells expressing mRuby3 alone, 16.5 ± 2.5% of the cells were FRET positive for tau aggregates, while cell lines expressing DNAJB6b_WT_-mRuby3 or DNAJB8_WT_-mRuby3 lead to a statistically significant 2-fold reduction in signal compared to mRuby3 alone (Figure 5B). We compared the %FRET_Clover/Cerulean_ in the cell populations expressing DNAJB8 mutants and reported the results normalized to DNAJB8_WT_ (Figure 5C). All seeded tau biosensor cell lines expressing DNAJB8 mutants yielded small increases in seeding compared to DNAJB8_WT_ but were not statistically significant, which suggests these mutations are not important for DNAJB8 holdase activity on tau. Cell viability was comparable across all DNAJB8-expressing tau biosensor lines including controls (Figure S5C), thus confirming that the processes were not affected by toxicity in response to co-expression of tau and DNAJB8. Consistent with our *in vitro* results, the designed DNAJB8 mutants that are monomeric retain activity similar to DNAJB8_WT_.

Using the mRuby3 tag on our DNAJB8 mutants, we performed an image-based analysis on each tau biosensor cell line to determine if the frequency of DNAJB8 puncta has changed in the presence of tau as described in Figure 1. First, we quantified mRuby3 signal across all lines transfected with a mRuby3-fusion construct to look at baseline expression. We observed similar variations in expression regardless of seeding, where relative to DNAJB8_WT_, the only mutants that showed a significant increase in mRuby3 signal per population were DNAJB8_S2_, DNAJB8_S3_, DNAJB8_S4_, DNAJB8_F148S_, and DNAJB8_F148S_F151S_ (Figure S5D). Overall, the average size of puncta and signal intensity was consistent across all cell lines (Figure S5E-F). Control cell lines expressing mRuby3 alone under unseeded and seeded with 20nM tau fibrils conditions yielded low frequency of puncta (Figure 5D; red). Expression of DNAJB6b_WT_-mRuby3 yielded 2-fold higher frequencies of puncta in unseeded and seeded conditions, and DNAJB8_WT_-mRuby3 yielded 6-fold higher puncta frequency compared to the control (Figure 5D). Under these conditions, DNAJB8 has a higher propensity to form large assemblies in cells compared to DNAJB6b. When we compared our DNAJB8 variants to the puncta frequency observed in DNAJB8_WT_-mRuby3 expressing tau biosensor cells, we observed the same trend as our HEK293 cells where mutations reduced puncta frequency for all mutants, with DNAJB8_S3_ reaching the same frequency as the mRuby3 control (Figure 5E). Our results suggest that the oligomeric assembly propensity of these two oligomeric JDPs is not related to their protein aggregation function, as supported by our data with mutant DNAJB8.

Finally, we asked whether our results on tau aggregation suppression by DNAJB8 in cells would translate to the well-established Htt substrate (Hageman, Rujano et al. 2010, Gillis, Schipper-Krom et al. 2013, Månsson, Kakkar et al. 2014, Kakkar, Mansson et al. 2016). We tested DNAJB8_WT_ and mutants on their ability to prevent aggregation of amyloidogenic fragment of the Huntingtin gene with extended polyglutamine chain (HttQ119) in cells using a Filter trap assay (FTA). HEK293T cells were co-transfected with a 1:9 ratio of Htt Q119-GFP: DNAJB8-mRuby3 and the amount of Htt aggregates were quantified by FTA (Figure 5F).

Both the expression of Htt Q119-GFP and the expression of DNAJB8-mRuby3 in DNAJB8_WT_ and all mutant expressing lines were confirmed to be similar by western blot (Figure S5G). Htt aggregate band intensity in the FTA was quantified and normalized to the mRuby3 negative control and reported as a percentage of aggregates of Q119 in this condition. As reported previously, DNAJB8_WT_ is a potent suppressor of Q119 aggregation yielding no signal (Figure 5G and Figure S5H). Similarly, little signal was detected in cells expressing DNAJB8_F148S_, DNAJB8_F151S_, DNAJB8_F148S_F151S_, and DNAJB8_S1_ (Figure 5G and Figure S5L). A minor loss in activity was found for DNAJB8_S2_, DNAJB8_S4_, and DNAJB8_S3_ (Figure 5G and Figure S5H), suggesting that most activity is retained even though some DNAJB8 mutants are unable to self-assemble. To ensure that the observed anti-aggregation activity is not influenced by the presence of endogenous DNAJB6b, we repeated the experiment using a HEK293T cell line in which DNAJB6b was knocked out (Thiruvalluvan, de Mattos et al. 2020). Also here, all DNAJB8 variants were able to reduce polyQ aggregation with DNAJB8_S2_, DNAJB8_S3_, and DNAJB8_S4_ relatively being slightly more active than in cells that did express endogenous DNAJB6b (Figure S5I-J).

The minor loss of activity of DNAJB8_S2_, DNAJB8_S4_ and DNAJB8_S3_ is not the result of changes in the oligomeric assembly of DNAJB8 since the monomeric minimal mutants DNAJB8_F151_ and DNAJB8_F148S_F151S_ retain equal chaperone activity relative to DNAJB8_WT_. Therefore, this loss of activity in DNAJB8_S3_ must be attributed to the loss of hydrophobic residues in the S/T-rich domain which seem more important for Htt activity compared to tau. Our combined data suggest that oligomeric assembly of DNAJB8 is not essential for substrate activity.

## DISCUSSION

The capacity of DNAJB6 and DNAJB8 to oligomerize has been established *in vitro* and in cells (Ryder, Matlahov et al. 2021),(Hageman, Rujano et al. 2010, Gillis, Schipper-Krom et al. 2013),(McMahon, Bergink et al. 2021). However, their mechanism of assembly has been poorly understood and remained intertwined with substrate activity. Previous studies suggested that DNAJB6 assembly may be driven by sequences along the C-terminus of the protein, similar to dimerization observed in other JDPs (Karamanos, Tugarinov and Clore 2019, Karamanos, Tugarinov and Clore 2020). This is not true for DNAJB8, where the CTD domain is a stable monomer that is unable to assemble (Ryder, Matlahov et al. 2021). Instead, DNAJB8 oligomerization was abolished by mutating aromatic residues in the disordered G/F-rich and S/T-rich domains (Ryder, Matlahov et al. 2021). By combining cellular and *in vitro* studies, we identified a small phenylalanine-rich region (i.e. S3) within the S/T-rich domain that is responsible for DNAJB8 self-assembly. We obtained an X-ray structure of the minimal assembly motif and by using mutagenesis verified that this motif (^147^AFSSFN^152^) is important for assembly of FL DNAJB8 *in vitro* and in cells. F→S mutations to positions in this motif yield monomeric DNAJB8 mutants *in vitro* with a more exposed S/T domain that retains activity against substrate aggregation *in vitro* and in cells. Our data builds a new paradigm for oligomerization of JDPs and for the first time decouples oligomerization from substrate activity underscoring that oligomerization is not a prerequisite for holdase activity. The S/T domain of oligomeric JDPs has been implicated in substrate binding activity, but as we discovered, other regions may play important specific substrate conformation and sequence dependent roles.

### Molecular interactions that drive DNAJB8 assembly

An emerging concept in the field of biology is the versatile way in which weak molecular nonpolar or charge complementary interactions can drive reversible (and irreversible) protein assembly to control spatial organization of compartments in cells. Proteins that encode low complexity domains, such as FUS, TDP-43, or HNRP1, use charge complementary or nonpolar residues to drive weak intermolecular interactions that promote phase separation, which result in dynamic and reversible assemblies (Pak, Kosno et al. 2016, Vernon, Chong et al. 2018, Dignon, Best and Mittal 2020, Krainer, Welsh et al. 2021). In particular, aromatic-aromatic residue interactions are especially stabilizing through π-π interactions (McGaughey, Gagne and Rappe 1998, Vernon, Chong et al. 2018). Intrinsically disordered proteins such as tau, aβ, or α-synuclein have been shown to initiate self-assembly through the formation of steric zippers, which are mediated by a combination of side chain and backbone hydrogen bonds (Sawaya, Sambashivan et al. 2007, Hughes, Sawaya et al. 2018). In addition, the role of backbone interactions in phase separation is unclear, but recent studies have suggested that a combination of backbone and side chain interactions may be critical for phase separation of low-complexity sequences (Zhou, Sumrow et al. 2022). Our data support that DNAJB8 oligomerization is dependent on both the side chain interactions as well β-strand mediated backbone hydrogen bonds (Figure 6). The combination of both of these intermolecular forces promotes the initial dimer assembly, followed by the hydrophobic interactions that yield larger oligomeric assemblies. We demonstrate that in both peptides and FL DNAJB8 disruptions to either the side chains (F→S) or the backbone (S→P) result in a loss of oligomeric assembly, indicating that a combination of hydrophobic “stickers” and backbone hydrogen bonds are equally important for DNAJB8 assembly (Figure 6). While low-complexity sequence motifs in other proteins may not necessarily share the same amino acid identities as DNAJB8, our findings present the idea that similar combinations of forces may be required for controlling reversible large-scale oligomeric assembly of a chaperone in cells. It is perhaps surprising that a chaperone would encode the capacity to adopt reversible assemblies, but one possible explanation is that the assemblies may be a storage conformation that persists in the cytosol, which can harbor monomers for future release to provide active chaperone conformations on demand.

**Figure 6.**
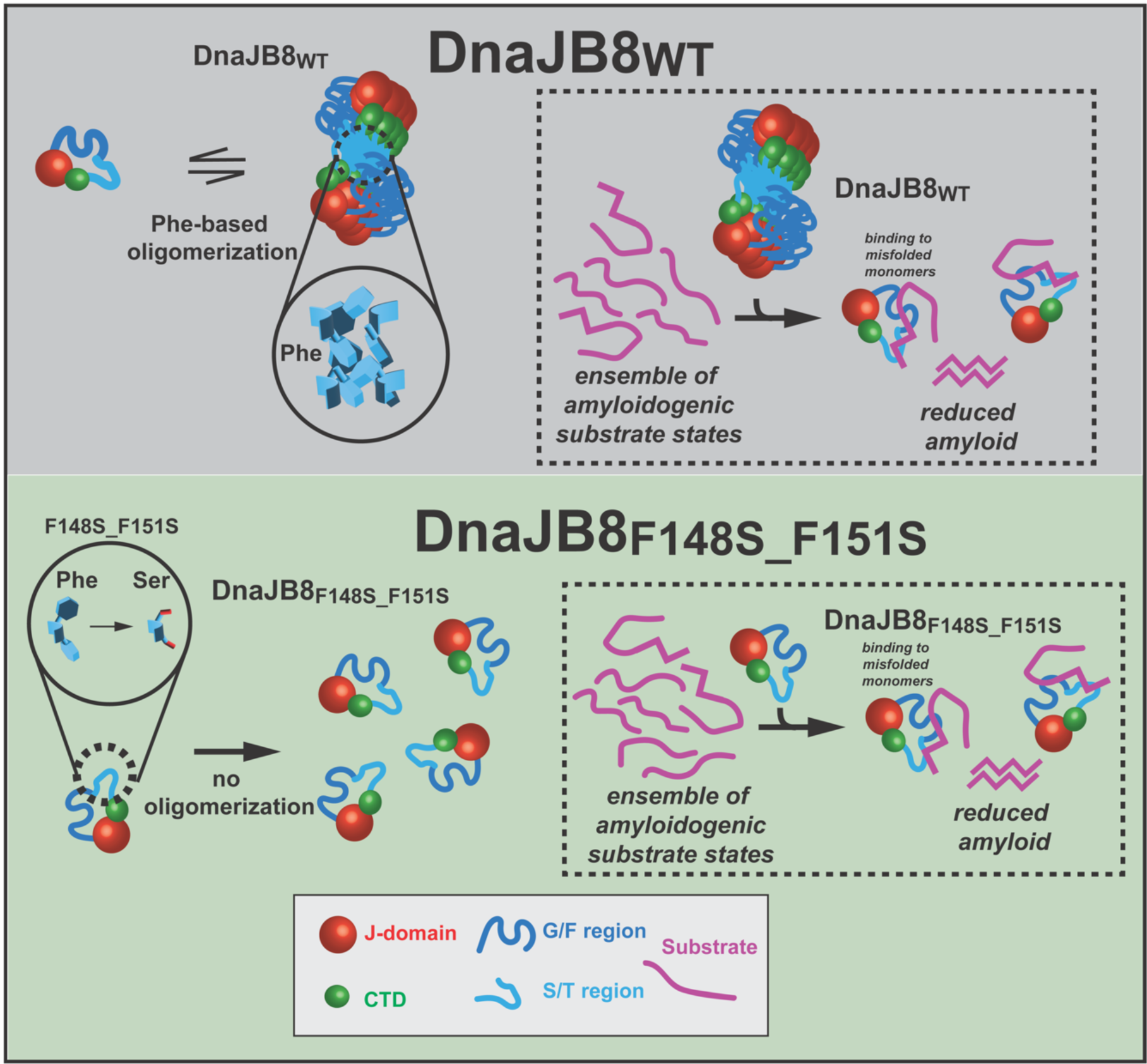
DNAJB8 oligomerization is not a prerequisite for activity. Model of DNAJB8 oligomeric assembly. The ^147^AFSSFN^152^ motif within the S/T-rich domain forms the core of the DNAJB8 oligomer with β-stranded dimers and Phe-Phe interactions stabilizing the hydrophobic core of the oligomer (top). Model showing how Phe→Ser mutations in the S/T-rich domain result in a mutant of DNAJB8 that is incapable of self-assembling. This results in DNAJB8 remaining a stable monomer in solution in the absence of Phe-Phe interactions (bottom). Model summarizing that DNAJB8 chaperone activity is independent on whether or not DNAJB8 exists as a monomer or oligomer (top right). DNAJB8 mutants where oligomerization was inhibited showed equal activity to DNAJB8_WT_ and other mutants that were still capable of forming oligomers in cells (top bottom). Therefore, the role of DNAJB8 oligomeric assemblies in cells is unrelated to its activity as a JDP chaperone.

### Oligomeric DNAJB8 interactions with substrate are dynamic

Chaperones often bind to substrates with micromolar affinities, which by design promotes substrate release and refolding (Hartl 1996, Kampinga and Craig 2010). This is true for large molecular chaperones such as chaperonins, Hsp90, as well as many JDPs (Qiu, Shao et al. 2006, Johnson 2012, Schopf, Biebl and Buchner 2017, Zarouchlioti, Parfitt et al. 2017). For globular substrates, chaperones interact with unfolded intermediates with weak affinities to prevent aggregation and allow refolding upon release into globular functional states. For intrinsically disordered proteins that do not adopt a defined globular fold, how do molecular chaperones regulate their folding? Our lab has uncovered that DNAJC7 binds tightly and specifically to the inert aggregation-resistant conformation of tau with nanomolar affinity (Hou, Wydorski et al. 2021). How oligomeric JDPs interact with substrates has remained an open question. The S/T-rich domain in oligomeric JDPs has been implicated in interactions with HttEx1and amyloidβ(Kakkar, Mansson et al. 2016, Osterlund, Lundqvist et al. 2020). Studying how oligomeric JDPs binding to substrates has been hampered by measuring interactions between polydisperse species of chaperone with substrates. This is further confounded by the proposed role of JDPs to interact with misfolded conformations of substrates. In this study we have uncoupled oligomerization of DNAJB8 from substrate activity to create a monomeric form of DNAJB8 that retains activity (Figure 6). Our methods to produce recombinant seeds of the tau protein (Mirbaha, Chen et al. 2018, Hou, Chen et al. 2021) allowed us to probe whether DNAJB8 mutants have differential activity against normal aggregation-resistant tau compared to a pathogenic monomer seed. Measuring affinities between the monomeric chaperone and these two forms of tau yield dramatic differences in binding affinity consistent with the chaperone preferentially interacting with misfolded seeds (Kakkar, Mansson et al. 2016).

Crosslinking of the mutant JDP to the two different conformations of tau reveals different patterns of interaction, and validates that the seed and normal tau differ in their shape and thus present different surfaces for JDP binding. Perhaps the largest difference is that the chaperone appears to bind more favorably with the basic PRD domain of tau, which may be stabilized by cation-π interactions. This is consistent with our prior work that the N-terminus rearranges its interaction with the repeat domain in the seeding form compared to M_i_(Chen, Drombosky et al. 2019). Based on the presently described π-π interactions that drive DNAJB8 oligomerization, it could be possible that large, positively charged surfaces along the tau PRD could interact with phenylalanine rings in a similar manner to cation-π interactions, resulting in a weak-charge based interaction between DNAJB8 and tau. In turn, this minor interaction would promote the extension of the DNAJB8 monomer and disengage the JD from the CTD in preparation for Hsp70 binding.

Despite the low affinities the monomeric JDP retains substrate aggregation prevention activity *in vitro* and in cells. When considering our activity data with HttEx1 Q119 in cells, the activity differences observed with DNAJB8_S3_ are likely due to the specific substitutions of five Phe positions as opposed to whether or not the protein is capable of self-assembly. This is consistent with previously reported data, where it has been proposed that the S/T-rich domain plays an important role in substrate recognition (Hageman, Rujano et al. 2010, Kakkar, Mansson et al. 2016). This leads to new questions about why the presence of Phe residues in the S/T-rich domain is important for DNAJB8 recognition of HttEx1, but the absence of those same residues improves *in vitro* recognition of tau. One possible explanation may rely upon previous work suggesting that the tau PRD is a common recognition domain to JDPs (Mok, Condello et al. 2018). This data for the first time uncovers differences in substrate activity and suggests that different surfaces on the JDP may play a role in substrate recognition. These data also raises another question: how does the chaperone influence aggregation of a substrate through low affinity interactions? Intrinsically disordered proteins sample different conformations and this is certainly true for the M_i_ conformation of tau. Misfolded seeds (i.e. M_s_) on the other hand adopts more defined alternate conformations. Based on previous experiments, we proposed that M_s_ preferentially exposes amyloid motifs (Mirbaha, Chen et al. 2018, Chen, Drombosky et al. 2019) and more recently we showed that the N-terminus of tau folds differently on the basic repeat domain (Hou, Chen et al. 2021). The higher binding affinity between the JDP and M_s_ suggest that the flexibility of normal inert tau lends to low affinities while the more defined and ordered M_s_ conformation leads to higher affinity possibly gained through PRD and amyloid motif interactions. Higher resolution structural experiments between engineered JDPs and substrates could uncover the details of these interactions.

### Substrate selectivity by oligomeric JDPs

DNAJB6 is ubiquitously expressed, including in the human brain, but its expression decreases with age (Hageman, Rujano et al. 2010, Kakkar, Mansson et al. 2016, Thiruvalluvan, de Mattos et al. 2020). DNAJB8 expression on the other hand is limited to the testes (Nishizawa, Hirohashi et al. 2012). Thus, the link between these JDPs and neurodegenerative diseases is weak because the onset of symptoms manifests later in life. Mutations in DNAJB6 are linked to muscular dystrophy, but the disease age onset varies (Sarparanta, Jonson et al. 2012). Initial genetic screens discovered these JDPs are regulators of Htt aggregation (Hageman, Rujano et al. 2010, Gillis, Schipper-Krom et al. 2013, Månsson, Kakkar et al. 2014, Kakkar, Mansson et al. 2016) and subsequent studies demonstrated their anti-aggregation activity against other IDPs (Aprile, Kallstig et al. 2017, Osterlund, Lundqvist et al. 2020, Thiruvalluvan, de Mattos et al. 2020, Arkan, Ljungberg et al. 2021, McMahon, Bergink et al. 2021). Why are these JDPs not expressed later in life when neurodegenerative diseases typically develop? Perhaps they only function at early stages of aggregation or alternatively their expression later in life has deleterious effects on other processes. What is the role of these JDPs? New evidence suggests that both DNAJB6 and DNAJB8 interact with FG nucleoporins to help assemble nuclear pores (Kuiper, Gallardo et al. 2022, Prophet, Rampello et al. 2022). FG Nups and IDPs belong to functionally different molecules and encode distinct sequence features. It is possible that these JDPs function on these two different classes of substrates using different folding mechanisms. Understanding in more detail how they function on substrates may allow us to engineer the different properties to increase their specificity, perhaps allowing them to be developed as therapeutics targeting specific proteins. Our study highlights that engineering DNAJB8 into a monomer allows it to maintain substrate activity but begins to change specificity between tau and Htt suggesting that distinct sequence features of the chaperone are used for different substrates. Future studies must be focused on understanding the role of this family of JDPs in maintaining cellular homeostasis, defining a clearer substrate interactome and understanding the mechanism of activity against substrates with different properties.

## METHODS

### Cell biological analysis of DNAJB8-mRuby3 lines

All plasmids for cell experiments were prepared by Twist Bioscience. The coding sequences for human DNAJB8_WT_, and the mutants DNAJB8_S1_, DNAJB8_S2_, DNAJB8_S3_, DNAJB8_S4_, DNAJB8_F148S_, DNAJB8_F151S_, and DNAJB8_F148S_F151S_ were cloned into FM5 lentiviral expression plasmids, in which the UbC promoter was replaced with a CMV promoter, the linker sequence was replaced by “GSAGSAAGSGEF,” and the YFP was replaced by mRuby3. The resulting genes produced a DNAJB8-mRuby3 fusion protein along with all mutant variants. In parallel, we produced a construct that expresses the fluorescent protein (mRuby3) but lacks any DNAJB8 gene. All plasmids were separately co-transfected into HEK293T cells using lipofectamine 2000 (Invitrogen). 100,000 cells were plated per well in media (10% FBS, 1% Pen/Strep, 1% GlutaMax in Dulbecco’s modified Eagle’s medium) in a 24-well glass bottom plate (Cellvis, P24-1.5-N). After 48h, cells were stained with Hoescht33342 at a final concentration of 2 μg/mL in cell media for 30 min at 37 °C and 5% CO_2_. The plate was placed on an IN Cell 6000 Analyzer (GE Healthcare) with a heated stage and 50 fields of view were imaged under 4′,6-diamidino-2-phenylindole (DAPI) and TxRed channels at ×60 magnification (Nikon ×60/0.95, Plan Apo, Corr Collar 0.11–0.23, CFI/60 lambda). Images were exported as TIFF files for downstream analysis. DNAJB8_WT_-mRuby3, all DNAJB8-mRuby3 mutants, mRuby3 alone, and HEK293T cells were plated and imaged in duplicates. Total cell counting, total mRuby3 expression, and total puncta counting was done using the CellProfiler v4.2.1 software.

### Determining oligomerization of DNAJB8 mutants by native gel protein electrophoresis

5×10^5^ HEK293 cells were plated on a 35 mm cell culture dish (Nunc) one day before transfection. Cells were transfected with 1 ug of plasmid DNA encoding the WT or mutant variants of DNAJB8 using polyethyleneimine (PEI) in a 1:3 ratio and incubated for 48 hours. After, cells were washed twice with cold phosphate-buffer saline (PBS), collected in Eppendorf tubes, and centrifuged at 500xg for 5 min. Cell pellets were lysed in a mild lysis buffer to preserve oligomeric structures (50 mM TRIS-HCl pH 7.4, 100 mM NaCl, 1 mM MgCl2, 0.5% NP-40, EDTA-free complete protease inhibitors cocktail (Roche), and 50 units/ml Denarase (c-LEcta)) on ice for 30 minutes with occasional (every 10 minutes) pipetting during the incubation period. The lysates were centrifuged at 14000 g for 10 min at 4° C, and the soluble fraction (supernatant) was used for further analysis. Protein concentration was measured with DC protein assay (Bio-Rad) and equalized using dilution buffer (50 mM TRIS-HCl pH 7.4, 100 mM NaCl, 1 mM MgCl2, 0.5% NP-40). Equalized protein lysates were mixed with non-denaturing sample buffer (4x, 125 mM Tris-HCl pH 6.7, 50% glycerol, 0.05% bromophenol blue sodium salt) and stored at -20° C.

For the native gel protein electrophoresis, samples were loaded on Mini-PROTEAN TGX Precast gel (4-15%, Bio-Rad) and ran in non-denaturing running buffer (25 mM Tris-HCl, 192 mM Glycine, pH 6.5) at 50V for the first hour with subsequent increase of the voltage to 100-110V. The protein electrophoresis system was placed in an ice bucket to maintain a low temperature (< 10° C) during the run; total running time - 7 hours. Proteins were transferred to nitrocellulose membrane (Schleicher and Schuell, PerkinElmer, Waltham, MA, USA) using Trans-Blot® Turbo™ Transfer System (Bio-Rad), blocked in 10% non-fat milk, and blotted with primary anti-DNAJB8 antibody (same as previously) overnight at 4° C on a rocking platform. After, the membrane was incubated with Anti-mouse HPR-conjugated secondary antibody (1:5000 dilution, GE Healthcare) at RT for 2 hours and visualized with enhanced chemiluminescence using ChemiDoc Imaging System (Bio-Rad).

### Recombinant DNAJB8 expression and purification

DNAJB8_WT_ and all DNAJB8 mutants were cloned into pET-29b (+) vectors and prepared by Twist Bioscience. DNAJB8-containing plasmids were transformed into *E.coli* BL21(DE3) competent cells, streaked, plated, and inoculated into 1L 2xLB, 0.05 mg/mL kanamycin and incubated at 37°C shaking at 220 rpm. Once OD_600_=0.6-0.8 AU, expression was induced upon adding 1mL of 1M IPTG to the 1L culture, and incubation continued for 4 hours at 37°C shaking at 220 rpm. Cells were harvested by spinning down the culture at 4,000xg for 20 min. The media was discarded and the cells were resuspended in 50mL lysis buffer (8M guanidinium HCl, 50mM HEPES, 20mM imidazole, 1mM DTT, pH 7.5). The cells were sonicated at 30% power, 5x pulse, for 10 min using an Omni Sonic Ruptor 4000 (Omni International). The lysed cells were pelleted at 10,000xg for 30 min and supernatant was retained. The lysate was mixed with 2mL HisPur™ Ni-NTA Resin (Thermo Scientific) for 1 hour before being loaded onto a gravity column. The column was washed with an additional 50mL of lysis buffer followed by 50mL of 50mM HEPES, 20mM imidazole, 1mM DTT, pH 7.5. The protein was eluted with 30mL elution buffer (50mM HEPES, 500mM imidazole, 1mM DTT, pH 7.5) and collected in 2mL fractions. High purity DNAJB8-containing fractions were pooled and loaded into 3.5 kDa Biotech CE Dialysis Tubing (Spectrum Labs) to be dialyzed overnight in 50mM ammonium formate at 4°C to minimize assembly formation while removing the imidazole. DNAJB8 was then lyophilized and stored at -80°C for future use.

### Dynamic light scattering

All DNAJB8 samples were prepared at 1.2 mg/mL in 1× PBS, 1 mM DTT pH 7.4. All protein samples were filtered through a 0.22-μm cellulose acetate sterile filter and loaded in triplicate onto a 384-well clear flat-bottom plate. The plate was loaded into a Wyatt DynaPro Plate Reader III and set to run continuously at room temperature at a scanning rate of 1 scan/15 min, with 1 scan composed of ten acquisitions. The data were analyzed using the Wyatt Dynamics software version 7.8.2.18. Light-scattering results were filtered by the sum of squares (SOS) < 20 to eliminate statistical outlier acquisitions within each scan. R_h_ of observed particles for two time points (0 and 30 h) was reported as histograms as a function of mass% for DNAJB8_WT_ and all DNAJB8 mutants. DNAJB8 samples were prepared at 10μM and tau M_s_ was prepared at 0.1μM for M_s_ assembly experiments. Data for M_s_ assembly experiments were plotted as average R_h_ of total particles over time. Data was not collected for some time points of the M_s_ experiment due to high SOS values and heterogeneity. It is expected during the aggregation phase of M_s_ that assembling fibrils exist at varying sizes until the population reaches equilibrium.

### SEC-MALS

DNAJB8_S3_, and DNAJB8_F148S_F151S_ constructs at a concentration of 4.2 mg/mL and 2.8 mg/mL, respectively, in 1xPBS were filtered through a 0.1 μm filter to remove larger impurities. Each sample was further filtered using a 0.22 μm centrifugal filter before 100 μL was applied to a Superdex 200 Increase 10/300 column equilibrated in 1xPBS with 1mM TCEP. The column was in line with a Shimadzu UV detector, a Wyatt TREOS II light-scattering detector, and a Wyatt Optilab tREX differential-refractive-index detector. The flow rate was 0.5 mL/min. The data were analyzed with Wyatt’s ASTRA software version 7.1.0.29. SEDFIT (Schuck 2000) was used to calculate the dn/dc of the protein.

### Thioflavin T fluorescence aggregation assays

For all endpoint assays, DNAJB8_WT_ and all DNAJB8 mutants were resuspended in 1xPBS, 1 mM DTT pH 7.4 to a final concentration of 50μM and incubated at room temperature for 30h. All peptides were analyzed at a concentration of 100μM. All samples were mixed with 25μM ThT and plated into a 384-well clear flat bottom plate in triplicate as technical replicates. ThT fluorescence was measured using a Tecan M1000 plate reader at 446 nm Ex (5 nm bandwidth), 482 nm Em (5 nm bandwidth). For peptide time course experiments, all peptides were suspended in a 1:1 mixture (v/v) of TFA (Pierce) and incubated at room temperature (RT) for 1h to monomerize all peptides. In a chemical fume hood, the peptide solution was dried under a stream of nitrogen gas, and then immediately placed under vacuum to remove any residual volatile solvents. The peptide residue was resuspended in 1xPBS, 1mM DTT and the pH restored to pH 7 using NaOH with a final concentration of 100μM. All peptides were mixed with 25μM ThT and plated into a 384-well clear flat bottom plate in triplicate as technical replicates. ThT fluorescence was measured using a Tecan M1000 plate reader at 446 nm Ex (5 nm bandwidth), 482 nm Em (5 nm bandwidth). For the tau aggregation experiment, DNAJB8 was resuspended in 1xPBS, 1 mM DTT pH 7.4 to a final concentration of 10μM, tau M_s_ was diluted in1xPBS, 1 mM DTT pH 7.4 to a final concentration of 0.05μM. tauRD was diluted in 1xPBS, 1 mM DTT pH 7.4 to a final concentration of 17μM followed by boiling at 100 °C for 5 min before being diluted further in 1xPBS, 1 mM DTT pH 7.4 to a final concentration of 5μM. 25μM ThT was mixed with all samples before the addition of M_s_ to all M_s_-containing reactions. All reactions were plated into a 384-well clear flat bottom plate in triplicate as technical replicates. ThT fluorescence was measured using a Tecan M1000 plate reader at 446 nm Ex (5 nm bandwidth), 482 nm Em (5 nm bandwidth).

### Transmission electron microscopy

Peptide samples were resuspended in 1xPBS, 1 mM DTT pH 7.4 to a final concentration of 100μM. 5 μL of sample was loaded onto a glow discharged Formvar-coated 300-mesh copper grid for 30 s and was blotted by filter paper followed by washing the grid with 5 μL of ddH2O. After another 30 seconds, 2% uranyl acetate was loaded on the grid and blotted again. The grid was dried for 1 min and loaded into a FEI Tecnai G2 Spirit Biotwin TEM. All images were captured using a Gatan 2K × 2K multiport readout post-column CCD at the UT Southwestern EM Core Facility.

### AFSSFN crystallization and structure determination

AFSSFN synthetic peptide (Genscript) was dissolved to 30 mg/mL in water and crystallized by hanging drop at a 1:2 ratio with 20% PEG 8000, 100 mM HEPES pH 7.5. X-ray diffraction data was collected at APS Beamline 24-ID-C. Diffraction data were indexed and scaled using XDS and XSCALE (Kabsch 2010). The SHELX macromolecular structure determination suite was used for phasing the measured intensities (Sheldrick 2008). Model-building and manual real-space refinement was performed in COOT (McCoy, Grosse-Kunstleve et al. 2007). Automated reciprocal-space and real-space refinement was performed using Refmac and Phenix (Murshudov, Vagin and Dodson 1997, Afonine, Grosse-Kunstleve et al. 2012). A summary of data collection and refinement statistics is given in Table S2.

### Tau Expression and Purification and Production of Seeding Monomer

Full-length recombinant tau was cloned into a pET28b vector as described previously (Chen, Drombosky et al. 2019). This vector was transformed into BL21(DE3) competent cells, and cultured at 37°C shaking at 220 rpm until OD_600_=1.4 AU. Expression was induced in each flask by the addition of 1mL 0.5M IPTG and continued incubation for 3.25 hours at 37°C shaking at 220 rpm. The cells were harvested by spinning down the cells at 5,000xg for 12 min and discarding the media. The cell pellets were combined and resuspended in 225mL lysis buffer (50mM Tris, 500mM NaCl, 20mM imidazole, 1mM βME, 1mM PMSF, pH 7.5) and sonicated at 30% power, 5x pulse, for 10 min on ice using an Omni Sonic Ruptor 4000 (Omni International). The lysate was spun down at 15,000xg for 20 min and the pellets were discarded. 6mL of HisPur™ Ni-NTA Resin (Thermo Scientific) was added to the lysate and incubated on a orbital shaker for 1 hour at 4°C. The lysate was loaded onto a gravity column and washed with 40CV of lysis buffer. The sample was eluted with 10mL (3CV) of elution buffer (50mM Tris, 250mM NaCl, 300mM imidazole, 1mM βME, pH 7.5) into 1mL fractions. Tau-containing fractions were pooled and desalted into 5mL 50mM MES, 50mM NaCl, 1mM βME, pH 6.0 using a PD10 desalting column containing Sephadex 25 resin (Cytiva). The sample was filtered and injected onto a 5×5mL HiTrap SP-HP cation exchange column (Cytiva) over a NaCl gradient of 50-200mM and collecting 2mL fractions for 80mL of elution volume. Tau-containing fractions were pooled and concentrated down to 500μL using a 10kDa MW cutoff Amicon Ultra-15 Centrifugal Filter (EMD Millipore). The sample was filtered and loaded onto a 10/300 Superdex 200 column (Cytiva) running either on 10mM HEPES, 100mM NaCl, 1mM DTT, pH 7.4 (Tau M_i_ elutions) or 30mM MOPS, 50mM KCl, 5mM MgCl_2_, 1mM DTT, pH 6.5 (Tau pre-M_s_ elutions).

Tau that was eluted into pre-M_s_ elution buffer was prepared in 2 tubes of 500μL at 8μM in pre-M_s_ elution buffer. One tube was injected onto a 10/300 Superdex 200 column (Cytiva) running 1xPBS, 1mM DTT, pH 7.4 to determine the elution profile for tau M_i_. The other tube was mixed with heparin at a 1:10 molar ratio (tau:heparin) and incubated at room temperature for 30 minutes to generate tau M_s_. The sample was then filtered and injected onto a 10/300 Superdex 200 column (Cytiva) running 1xPBS, 1mM DTT, pH 7.4, where fractions containing tau M_s_ were isolated from remaining tau M_i_ and oligomers. Samples were immediately flash frozen in liquid nitrogen and stored at -80°C for future experiments.

### Microscale thermophoresis

All MST experiments were performed on a Nanotemper Monolith NT.115 in the Molecular Biophysics Core at UTSW and analyzed with a standard protocol (Scheuermann, Padrick et al. 2016). All binding measurements were done as technical triplicates. 1μM tau M_i_ and M_s_ was labelled with 4mM Cy5-NHS by incubating in the dark for 30 minutes at room temperature. Excess dye was removed using a Microspin™ G-25 desalting column (Cytiva). Labelled tau was mixed with a 16-step titration of DNAJB8_S3_ between 0.006-100μM. Data were fit in PALMIST using a 1:1 binding model (Scheuermann, Padrick et al. 2016) and analyzed using GUSSI (Brautigam 2015).

### Crosslinking reagents

DMTMM (Sigma-Aldrich) is commercially available. For all cross-linking experiments, DMTMM (Sigma-Aldrich) was prepared at a 120 mg/mL concentration in 1× PBS pH 7 and stored at -80°C as a stock solution.

### Cross-linking mass spectrometry

For all DNAJB8 experiments, DNAJB8_WT_, DNAJB8_S1_, DNAJB8_S2_, DNAJB8_S3_, and DNAJB8_S4_ were all resuspended in 1xPBS, 1mM DTT pH 7.4 buffer to a final concentration of 25μM. For DNAJB8 with M_s_ experiments, DNAJB8_WT_ and DNAJB8_S3_ were resuspended in 1xPBS, 1mM DTT pH 7.4 buffer to a final concentration of 10μM and M_s_ was diluted in 1xPBS, 1mM DTT pH 7.4 buffer to a final concentration of 0.1μM. All experiments were performed in triplicate. All samples were incubated at 37 °C while shaking at 350 r.p.m. for 30 min. Final concentrations of 36mM DMTMM (Sigma-Aldrich) were added to the protein samples and incubated at 37 °C with shaking at 350 r.p.m. for 30 min. The reactions were quenched with 100mM ammonium bicarbonate and incubated at 37 °C for 30 min. Samples were lyophilized and resuspended in 8M urea. Samples were reduced with 2.5mM tris (2-carboxyethyl)phosphine (TCEP) incubated at 37 °C for 30 min, followed by alkylation with 5mM iodoacetimide for 30 min in the dark. Samples were diluted to 1M urea using a stock of 50mM ammonium bicarbonate and trypsin (Promega) was added at a 1:50 enzyme-tosubstrate ratio and incubated overnight at 37 °C while shaking at 600 r.p.m. Two percent (v/v) formic acid was added to acidify the samples following overnight digestion. All samples were run on reverse-phase Sep-Pak tC18 cartridges (Waters) eluted in 50% acetonitrile with 0.1% formic acid. Ten microliters of the purified peptide fractions were injected for liquid Chromatography with tandem mass spectrometry analysis on an Eksigent 1D-NanoLC-Ultra HPLC system coupled to a Thermo Orbitrap Fusion Tribrid System. The MS was operated in data-dependent mode by selecting the five most abundant precursor ions (m/z 350–1600, charge state 3+ and above) from a preview scan and subjecting them to collision-induced dissociation (normalized collision energy = 35%, 30 ms activation). Fragment ions were detected at low resolution in the linear ion trap. Dynamic exclusion was enabled (repeat count 1, exclusion duration 30 s).

### Analysis of MS results

All MS experiments were carried out on an Orbitrap Fusion Lumos Tribrid instrument available through the UTSW proteomics core facility. Each Thermo.raw file was converted to .mzXML format for analysis using an in-house installation of xQuest. Score thresholds were set through xProphet, which uses a target/decoy model. The search parameters were set as follows: maximum number of missed cleavages = 2, peptide length = 5–50 residues, fixed modifications carbamidomethyl-Cys (mass shift = 57.02146 Da), mass shift of cross-linker = −18.010595 Da, no monolink mass specified, MS1 tolerance = 15 p.p.m., and MS2 tolerance = 0.2 Da for common ions and 0.3 Da for cross-link ions; search in enumeration mode.

### Cell biological analysis of tau biosensor lines expressing DNAJB8-mRuby3

Original FM5 plasmids used for generating the tauRD (P301S) cells were previously described (Holmes, Furman et al. 2014, Sanders, Kaufman et al. 2014), and modified for these experiments as previously described (Hitt, Vaquer-Alicea et al. 2021). HEK293T cells were co-transfected with lentivirus encoding tauRD (P301S)-mClover3/mCerulean3 and sorted as previously described (Hitt, Vaquer-Alicea et al. 2021) to generate stable cell lines. All DNAJB8-containing plasmids for cell experiments were prepared by Twist Bioscience and constructed as described (see Cell biological analysis of DNAJB8-mRuby3 lines). tauRD (P301S) biosensor cells were plated at 10,000 cells/well in media (10% FBS, 1% Pen/Strep, 1% GlutaMax in Dulbecco’s modified Eagle’s medium) in a 96-well glass bottom plate (Cellvis, P96-1.5-N). 75ng DNAJB8-mRuby3 plasmids were separately co-transfected into the cells using lipofectamine 2000 (Invitrogen), with each DNAJB8 construct plated in 6 replicate wells. At 24 or 72 hours after DNAJB8-mRuby3 transfection, 20nM sonicated full-length tau fibrils were transfected into the cells using lipofectamine 2000 (Invitrogen). Tau fibrils were transfected into 3/6 wells per condition, resulting in triplicate wells for each DNAJB8 construct with and without tau seeding. After 24 hours, cells were stained with DRAQ5 at a final concentration of 5μM in cell media for 15 min at 37 °C and 5% CO_2_. The plate was loaded into an ImageXpress Confocal HT.ai (Molecular Dynamics) with a heated stage and 64 fields of view were taken using far red, FITC, CFP, and TxRed channels at 20x magnification (Nikon 20x/0.80, Plan Apo λD, WD 0.8, CS 0.17, MRD70270). Images were acquired in the MetaXpress software (Molecular Dynamics) and exported as TIFF files for downstream analysis. Total cell counting, total mClover3 expression, total mRuby3 expression, and total DNAJB8 puncta counting was done using the CellProfiler v4.2.1 software.

### Flow cytometry analysis of tau biosensor lines

The LSRFortessa SORP (BD Biosciences) was used to perform FRET flow cytometry on tau biosensor cells to measure aggregation. We used a gating strategy to isolate the cell population that expressed the fluorescent proteins, tau tagged to either mCerulean3 and mClover3 and DNAJB8-mRuby (Fig 5A). The gating strategy for this selection is highlighted in Fig S5C. Subsequently, within this population containing DNAJB8 expression, tau aggregation was calculated by measuring FRET of mCerulean3 bleed-through into the mClover3. Briefly, to measure mCerulean3 and FRET signal, cells were excited with the 405 nm laser and fluorescence was captured with a 405/50-nm and 525/50-nm filter, respectively. To measure mClover, cells were excited with a 488-nm laser and fluorescence was captured with a 525/50 nm filter. FRET channels were compensated using the BD FACSDiva Software, as described previously (Holmes, Furman et al. 2014, Furman, Holmes and Diamond 2015). FRET signal is defined as the percentage of FRET-positive cells in all analyses. The FRET population was adjusted with cells that received lipofectamine alone and are FRET-negative. In each experiment, at least 10,000 mCerulean3/mClover3 positive cells were analyzed as biological triplicates. Data analysis was performed using FlowJo v10 software (Treestar).

### Q119 cell culture and transient transfection

HEK293T were cultured at 37° C and 5% CO2 in DMEM (Gibco) supplemented with 10% FBS (Fetal bovine serum, Bodinco) and 100 U/ml penicillin/streptomycin (Invitrogen). For transient transfections, 5×10^5^ cells were plated on 35 mm cell culture dishes (Nunc) one the day before. Cells were transfected with 2 ug of total plasmid DNA (0.2 µg pQ119-GFP, 1.8 µg pDNAJB8 variant/empty vector) using polyethyleneimine (PEI) in 1:3 ratio.

### Protein extraction for FTA and western blot

24 h after transfection, cells were washed twice with cold phosphate-buffer saline (PBS), scraped in 230 µl lysis buffer (50 mM TRIS-HCl pH 7.4, 100 mM NaCl, 1 mM MgCl_2_, 0.5% SDS, EDTA free complete protease inhibitors cocktail (Roche), and 50 units/ml Denarase (c-LEcta)), and left on ice for 30 minutes with occasional vortexing. After incubation, SDS concentration was increased to 2%. Protein concentrations were measured with DC protein assay (Bio-Rad), equalized using dilution buffer (50 mM TRIS-HCl pH 7.4, 100 mM NaCl, 1 mM MgCl_2_, 2% SDS, 50 mM DTT), boiled for 5 minutes, and stores at -20° C.

### Filter trap assay

For filter trap assay (FTA), prepared samples were diluted 5-fold in dilution buffer. Then, both original (1x) and dilute (0.4x) samples were applied onto a 0.2 µm pore Cellulose acetate membrane, prewashed with FTA buffer (10 mM TRIS-HCl pH 8.0, 150 mM NaCl, 0.1% SDS). The membrane was washed under mild suction three times with FTA buffer, blocked in 10% non-fat milk, and blotted with anti-GFP/YFP antibody (mouse, Clontech) at 1:5000 dilution overnight at 4° C on a rocking platform. After, the membrane was incubated with an HPR-conjugated secondary antibody and visualized with enhanced chemiluminescence using ChemiDoc Imaging System (Bio-Rad). Signal intensities were measured in ImageLab Software, and the results of two dilutions (where possible) were averaged. Values were normalized to the control sample, analyzed with GraphPad Prism, and plotted in a graph. The resulting graph represents an average of three separate experiments.

### Western Blot analysis

For WB, samples aliquots were mixed with 4x Sample buffer (50 mM Tris-HCl pH 6.7, 2% SDS, 10% glycerol, 12.5 mM EDTA, 0.02% Bromophenol blue). Samples were loaded on 10% SDS-PAGE gel and ran at 90 V. Proteins were transferred to nitrocellulose membrane (Schleicher and Schuell, PerkinElmer, Waltham, MA, USA), blocked in 10% non-fat milk, and blotted with primary anti-GAPDH antibody (Sigma) diluted 1:5000, or with anti-DNAJB8 antibody (monoclonal, mouse, clone #EMR-DNAJB8.214-8) overnight at 4° C on a rocking platform. The anti-DNAJB8 antibody was prepared as previously described (Inoda, Hirohashi et al. 2009, Nishizawa, Hirohashi et al. 2012). After, the membrane was incubated with an HPR-conjugated secondary antibody and visualized with enhanced chemiluminescence using ChemiDoc Imaging System (Bio-Rad).

## ACKNOWLEDGEMENTS

L.A.J is supported by an Effie Marie Cain Scholarship in Medical Research and a grant from the NIH-NIA (R01AG065407). Mass spectrometry experiments were carried out at the UTSW Proteomics core. Transmission electron microscopy was performed at the Electron Microscopy Core Facility at UTSW, which is supported by the National Institutes of Health (NIH) (1S10OD021685-01A1 and 1S10OD020103-01). We would like to thank Dr. Andrew Lemoff from the UTSW Proteomics Core for his valuable insights on troubleshooting procedures in cross-linking mass spectrometry. We would like to thank Dr. Chad Brautigam from the Molecular Biophysics Resource for his guidance on SEC-MALS experiments. We want to thank Vaibhav Bommareddy for preparing the plasmids of CMV FM5 tau P301S. We thank Mike Collazo at the UCLA-DOE X-ray Crystallization and Crystallography Core Facilities, which are supported by DOE Grant DE-FC02-02ER63421. We thank M. Capel, K. Rajashankar, N.Sukumar, J.Schuermann, I. Kourinov and F. Murphy at NECAT beamlines 24-ID at APS, which are supported by grants from the National Center for Research Resources (5P41RR015301-10) and the National Institute of General Medical Sciences (8 P41 GM103403-10) from the National Institutes of Health. Use of the APS is supported by DOE under Contract DE-AC02-06CH11357. We thank all members of the Joachimiak lab for discussions and input on the manuscript.

## CONTRIBUTIONS

B.D.R. and L.A.J. initiated the project. B.D.R purified all proteins involved in the study. B.D.R. performed all peptide and protein assembly experiments. B.D.R performed all cell experiments and image analyses. B.D.R and A.O. performed the flow cytometry analysis. D.R.B. and M.R.S. determined the X-ray structure of the peptide. B.D.R and M.I.K. performed the gel crosslinking experiments. P.M. and N. M. performed the SEC-MALS experiments. P.M.W. performed the MST binding experiments. B.D.R performed the XL-MS analysis. E.U. performed native gel analysis of DNAJB8 samples and polyQ cell experiments. M.I.D and H.H.K. provided guidance on cell experiments. Finally, B.D.R and L.A.J. conceived of and directed the research as well as wrote the manuscript. All authors contributed to the revisions of the manuscript.

## DATA AVAILABILITY

The structural datasets for the AFSSFN peptide assembly generated during the current study are available in the Protein Data Bank repository (https://www.rcsb.org/) under PDB id: 8DTS. Source data are provided with this paper. Analysis of WT and mutant DNAJB8-mRuby3 expression in HEK293T cells and HEK293T cells expressing tauRD are available in Source Data 1. Raw DLS data are available in Source Data 2. Raw XL-MS data are available in Source Data 3. Raw data used in generating all other plots along with raw western blots and SDS gels are available in Source Data 4. All other data generated during the current study are available upon reasonable request.

## DECLARATION OF INTERESTS

The authors declare that they have no competing interests.

## Notes

### Competing Interest Statement

The authors have declared no competing interest.

### Summary of Updates

Updated version with changes for resubmission

