## Supplementary Information File for "DNAJB8 oligomerization is mediated by an aromatic-rich motif that is dispensable for substrate activity"

**Source Data 1. Raw data from cell analyses used in the study.**

**Source Data 2. Raw data from DLS data used in the study.**

**Source Data 3. Raw XL-MS data used in the study.**

**Source Data 4. Raw data from fluorescence assays, SASA calculations, FTA, SDS-PAGE, and western blots used in the study.**

### Supplemental Tables

| Protein Origin | Name | Peptide Sequence | Domain(s) |
|---|---|---|---|
| DNAJB8 | B8_1 | N(acetylation)-SLYDRAGCDSWRAGGGAST-C(amidation) | JD+G/F |
| DNAJB8 | B8_2 | N(acetylation)-PYHSPFDTGYTFRNPEDIFR-C(amidation) | G/F |
| DNAJB8 | B8_3 | N(acetylation)-EFFGGLDPFSFEFWDSPFNS-C(amidation) | G/F |
| DNAJB8 | B8_4 | N(acetylation)-DRGGRGHGLRGAFSAGFGEF-C(amidation) | G/F |
| DNAJB8 | B8_5 | N(acetylation)-PAFMEAFSSFNMLGCSGGSH-C(amidation) | G/F+S/T |
| DNAJB8 | B8_6 | N(acetylation)-TTFSSTSFGGSSSGSSGFKS-C(amidation) | S/T |
| DNAJB8 | B8_7 | N(acetylation)-VMSSTEMINGHKVTTKRIVE-C'(amidation) | CTD |
| DNAJB8 | B8_8 | N(acetylation)-NGQERVEVEEDGQLKSVTVN-C(amidation) | CTD |
| DNAJB8 | F148S | N(acetylation)-PAFMEASSSFNMLGCSGGSH-C(amidation) | G/F+S/T |
| DNAJB8 | F151S | N(acetylation)-PAFMEAFSSSNMLGCSGGSH-C(amidation) | G/F+S/T |
| DNAJB8 | F148S F151S | N(acetylation)-PAFMEASSSSNMLGCSGGSH-C(amidation) | G/F+S/T |
| DNAJB8 | S149P | N(acetylation)-PAFMEAFPSFNMLGCSGGSH-C(amidation) | G/F+S/T |
| DNAJB8 | S149P S150P | N(acetylation)-PAFMEAFPPFNMLGCSGGSH-C(amidation) | G/F+S/T |
| DNAJB6b | B6_5 | N(acetylation)-PSFGSGFSSFDTGFTSFGSLGHGGL-C(amidation) | G/F+S/T |
| DNAJB6b | B6_6 | N(acetylation)-TSFSSTSFGGSGMGNFKSISTST-C(amidation) | S/T |
| DNAJB6b | B6_7 | N(acetylation)-KMVNGRKITTKRIVENGQERVEVEED-C(amidation) | CTD |
| DNAJB6b | B6_8 | N(acetylation)-GQLKSLTINGKEQLLRLDNK-C(amidation) | CTD |

### Table S1. DNAJB8 peptides used for ThT and TEM screening

| | |
|---|---|
| X-ray Source | APS 24-ID-C |
| Wavelength | 0.77490 |
| Resolution Range | 11.92 – 0.75 (0.7769 – 0.75) |
| Space Group | P1 |
| Unit Cell | 38.38, 9.48, 30.4, 90, 90.001, 90.007 |
| Total Reflections | 182237 (8516) |
| Unique Reflections | 48629 (3777) |
| Multiplicity | 3.7 (2.3) |
| Completeness (%) | 88.28 (67.99) |
| Mean I/sigma(l) | 5.72 (1.32) |
| Wilson B-factor | 3.64 |
| R-merge | 0.1128 (0.5809) |
| R-meas | 0.1277 (0.7468) |
| R-pim | 0.05846 (0.4654) |
| CC1/2 | 0.99 (0.704) |
| CC* | 0.997 (0.909) |
| Reflections used in refinement | 48488 (3772) |
| Reflections used for R-free | 4851 (376) |
| R-work | 0.1989 (0.3118) |
| R-free | 0.2385 (0.3533) |
| CC(work) | 0.977 (0.868) |
| CC(free) | 0.964 (0.698) |
| Number of non-hydrogen atoms | 928 |

| | |
|---|---|
| Macromolecules | 886 |
| Solvent | 42 |
| Protein Residues | 72 |
| RMS(bonds) | 0.014 |
| RMS(angles) | 1.51 |
| Ramachandran Favored (%) | 100.00 |
| Ramachandran Allowed (%) | 0.00 |
| Ramachandran Outliers (%) | 0.00 |
| Rotamer Outliers (%) | 0.00 |
| Clashscore | 3.74 |
| Average B-factor | 14.76 |
| Macromolecules | 14.29 |
| Solvent | 24.60 |

**Table S2. Summary of X-ray data collection and refinement statistics**

**$^{147}$AFSSFN$^{152}$**

### Supplemental Figures and Legends

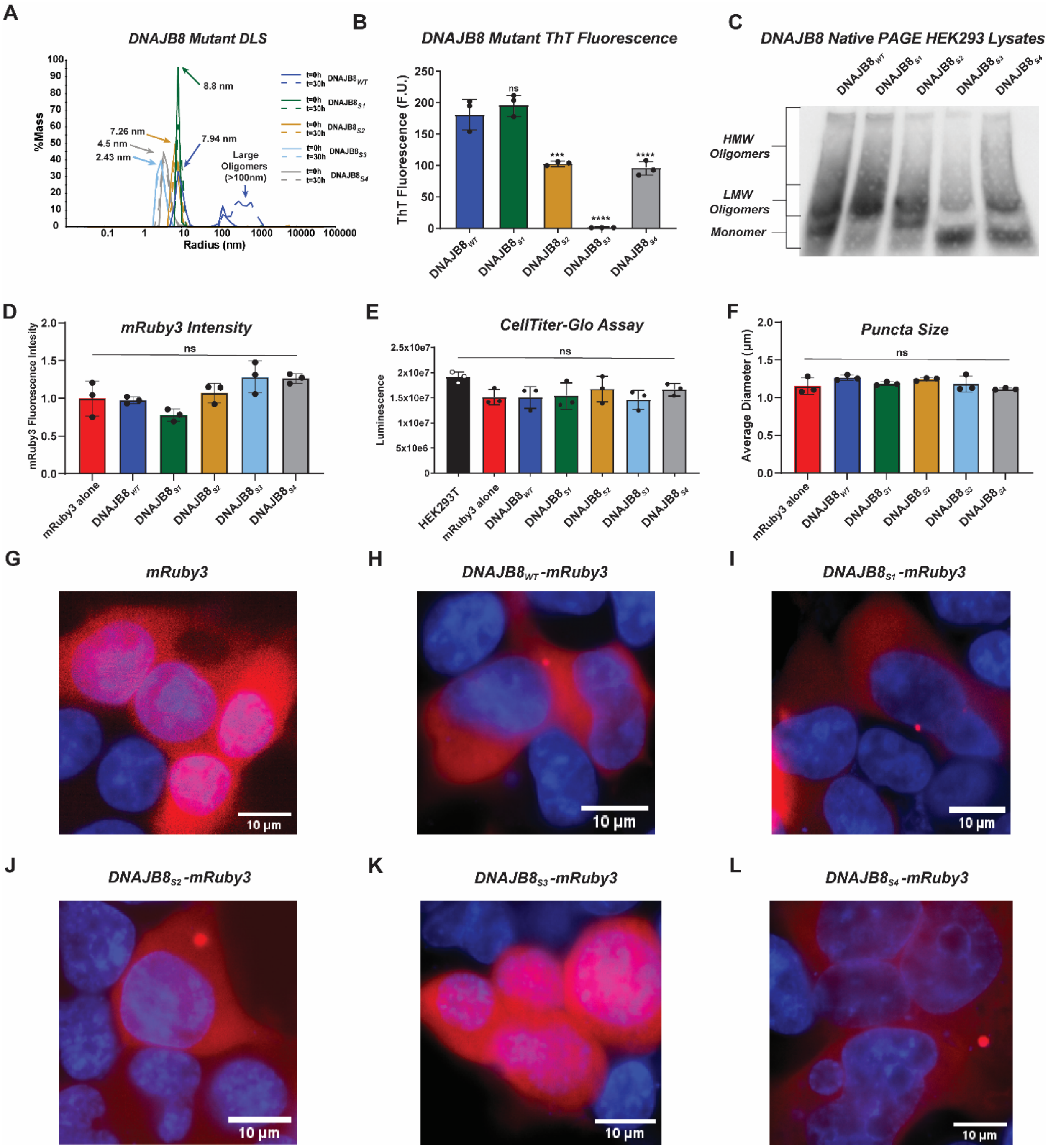


**Figure S1. Cell image analysis statistics for expression WT and segmented mutants of DNAJB8.** (**A**) DLS histogram reporting $R_h$ distributions of $DNAJB8_{WT}$ (blue), $DNAJB8_{S1}$ (green), $DNAJB8_{S2}$ (gold), $DNAJB8_{S3}$ (cyan), and $DNAJB8_{S4}$ (grey) at time points t=0hrs (solid lines) and t=30hrs (dashed lines). While mutations to all mutants resulted in slower assembly, DNAJB8$_{S1}$ and DNAJB8$_{S2}$ showed size distributions consistent with DNAJB8$_{WT}$, while DNAJB8$_{S3}$ and DNAJB8$_{S4}$ were more compact, consistent with stable monomers with buried hydrophobic residues. Shown values were calculated with SOS < 20. (**B**) Bar plot showing Thioflavin T endpoint fluorescence after 30h incubation at room temperature. Average fluorescence signal across triplicate technical replicates was reported for DNAJB8$_{WT}$ (blue), DNAJB8$_{S1}$ (green), DNAJB8$_{S2}$ (gold), DNAJB8$_{S3}$ (cyan), and DNAJB8$_{S4}$ (grey). DNAJB8$_{S1}$ fluorescence signal was comparable to self-assembly prone DNAJB8$_{WT}$, while DNAJB8$_{S2}$ and DNAJB8$_{S4}$ showed a ~2-fold reduction in fluorescence signal. DNAJB8$_{S3}$ was the only protein to remain ThT negative. One-way ANOVA was used to report significance where F= 101.2 and P-value<0.0001. For multiple comparisons to DNAJB8$_{WT}$: *** p<0.001, and **** p<0.0001. (**C**) Western blot of lysates from HEK293T cells expressing DNAJB8$_{WT}$-mRuby3, DNAJB8$_{S1}$-mRuby3, DNAJB8$_{S2}$-mRuby3, DNAJB8$_{S3}$-mRuby3, and DNAJB8$_{S4}$-mRuby3 run on native PAGE and treated with α-DNAJB8. Regions corresponding to monomers, low molecular weight oligomers, and high molecular weight oligomers are marked. (**D**) Transient expression of mRuby3 (red), DNAJB8$_{WT}$-mRuby3 (blue), DNAJB8$_{S1}$-mRuby3 (green), DNAJB8$_{S2}$-mRuby3 (gold), DNAJB8$_{S3}$-mRuby3 (cyan), and DNAJB8$_{S4}$-mRuby3 (grey) in HEK293T cells. Expression is quantified by averaging the signal intensities across all captured images normalized per the number of cells. These data were used to demonstrate that the results shown in Figure 1D are independent of protein overexpression variation. (**E**) CellTiter-Glo luminescence for each cell line as described above. The cell viability of each population is quantified through luminescence. These data demonstrate that the results shown in Figure 1D are independent of potential cell death. (**F**) Average diameter of puncta counted by CellProfiler across different cell lines. All cell line proteins are the same as in panel a. On average, most puncta detected in cells were around 1µm in diameter, regardless of mutations to DNAJB8. (**G**) Image of cells expressing mRuby3 (red) with nuclei stained with DAPI (blue). When expressed alone, mRuby3 signal is evenly and ubiquitously distributed across the cell with no localization bias. Scale bar= 10µm (**H**) Image of cells expressing DNAJB8$_{WT}$-mRuby3 (red) with nuclei stained with DAPI (blue). Expression of mRuby3-tagged DNAJB8$_{WT}$ is limited to the cytoplasm, where small inclusions can be observed. Scale bar= 10µm. (**I**) Image of cells expressing DNAJB8$_{S1}$-mRuby3 (red) with nuclei stained with DAPI (blue). Expression of mRuby3-tagged DNAJB8$_{S1}$ is limited to the cytoplasm, where small inclusions can be observed. Scale bar= 10µm. (**J**) Image of cells expressing DNAJB8$_{S2}$-mRuby3 (red) with nuclei stained with DAPI (blue). Expression of mRuby3-tagged DNAJB8$_{S2}$ is limited to the cytoplasm, where small inclusions can be observed. Scale bar= 10µm. (**K**) Image of cells expressing DNAJB8$_{S3}$-mRuby3 (red) with nuclei stained with DAPI (blue). Expression of mRuby3-tagged DNAJB8$_{S3}$ ubiquitous as the protein is now observed both in the cytoplasm and nucleus of the cell. Instances of inclusions are rare. Scale bar= 10µm. (**L**) Image of cells expressing DNAJB8$_{S4}$-mRuby3 (red) with nuclei stained with DAPI (blue). Expression of mRuby3-tagged DNAJB8$_{S4}$ is limited to the cytoplasm, where small inclusions can be observed. Scale bar= 10µm.

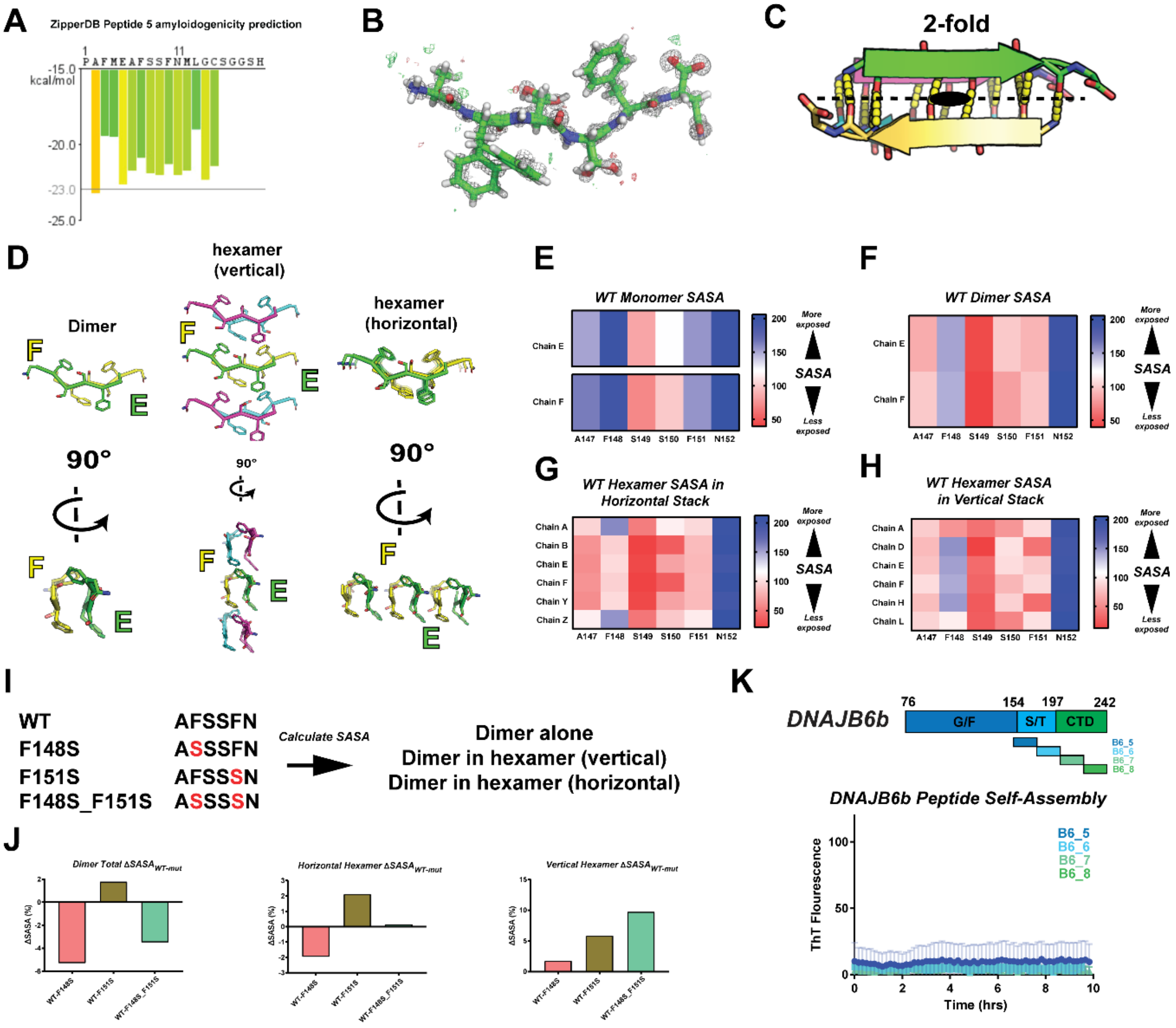


**Figure S2. Structural analysis of the $^{147}$AFSSFN$^{152}$ fragment.** (**A**) ZipperDB prediction of the B8_5 peptide fragment. Data is shown as a bar plot of six amino acid windows across the full fragment. The y-axis represents Rosetta Energy Units (REU). (**B**) $2F_o$-$F_c$ electron density map for the $^{147}$AFSSFN$^{152}$ fragment contoured at 1σ and $F_o$-$F_c$ is contoured at 3σ (green and red). $^{147}$AFSSFN$^{152}$ is shown in stick representation. (**C**) Illustration of the two-fold symmetry between two $^{147}$AFSSFN$^{152}$ anti-parallel strands. The structure is shown in cartoon representation. 2-fold axis is indicated by oval and dashed line. (**D**) Illustration of the interactions formed between two anti-parallel strands (left), vertical layers (middle) and horizontal layers (right). Structures are shown in stick representation. (**E**) SASA analysis of the $^{147}$AFSSFN$^{152}$ peptide monomers when loaded into Rosetta. SASA is calculated per reside in both chains E and F where blue indicates more solvent exposure, and red indicates greater burial within the structure. (**F**) SASA analysis of the $^{147}$AFSSFN$^{152}$ peptide dimer where chain E and F are paired together based on the structure from panel 2d. The interacting dimer pair results in greater burial within the core of the peptide dimer. (**G**) SASA analysis of the $^{147}$AFSSFN$^{152}$ peptide horizontal assembly using 6 chains of the peptide. As shown by the Rosetta model, this arrangement results in greater burial of F148 when the oligomer is constructed from the β-strand backbone. (**H**) SASA analysis of the $^{147}$AFSSFN$^{152}$ peptide vertical assembly using 3 peptide dimers stacked vertically. Here, F151 shows increased burial as this assembly is more dependent on the Phe-Phe stacking within the core of the oligomer. (**I**) Schematic for structural analysis of SASA changes in different interactions in the $^{147}$AFSSFN$^{152}$ assembly upon serine mutation at 148, 151 and 148_151. (**J**) Summary of ΔSASA shifts between the WT $^{147}$AFSSFN$^{152}$ dimer and oligomeric assemblies vs. Rosetta generated F148S, F151S, and F148S F151S mutant peptides. These results show that F151S results in increased SASA for all instances, which would imply this position is important in the Phe-Phe stacking arrangement of the DNAJB8 oligomer. (**K**) Thioflavin T assay using homologous S/T-rich domain and CTD peptide sequences from the homologous DNAJB6b protein. Unlike the DNAJB8-derived peptide B8_5, the homologous DNAJB6b sequence (B6_5) does not self-assemble into ThT positive structures, nor do any other homologous sequences within both domains. Thus, the ThT positive arrangement observed in DNAJB8 is specific to DNAJB8.

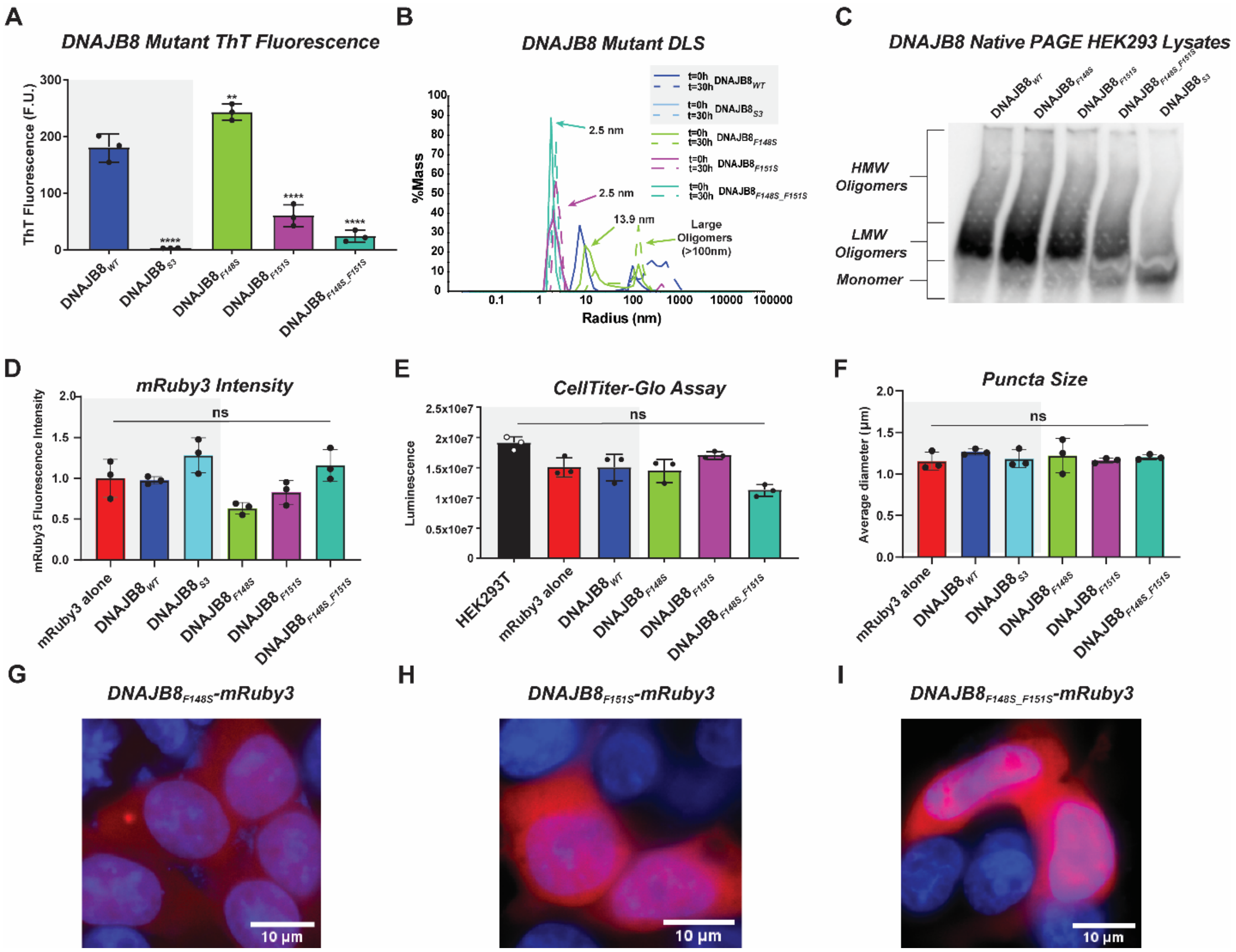


**Figure S3. Cell image analysis statistics for single and double substitution mutants at F148 and F151.** (**A**) Bar plot showing Thioflavin T endpoint fluorescence after 30 h incubation at room temperature. Average fluorescence signal across triplicate technical replicates was reported for $DNAJB8_{WT}$ (blue), $DNAJB8_{S3}$ (cyan), $DNAJB8_{F148S}$ (lime), $DNAJB8_{F151S}$ (purple) and $DNAJB8_{F148S\_F151S}$ (teal). Data for $DNAJB8_{WT}$ and $DNAJB8_{S3}$ (gray box) was previously shown in Figure 1D. While F148S in the context of FL DNAJB8 did not show any loss of self-assembly by ThT fluorescence compared to $DNAJB8_{WT}$, both $DNAJB8_{F148S}$ and $DNAJB8_{F148S\_F151S}$ showed considerably less ThT fluorescence, akin to decreased β-stranded self-assembly. Statistics calculated by one-way ANOVA. ** $p<0.01$, **** $p<0.0001$. (**B**) DLS histogram of $R_h$ distributions across $DNAJB8_{WT}$ (blue), $DNAJB8_{S3}$ (cyan), $DNAJB8_{F148S}$ (lime), $DNAJB8_{F151S}$ (purple) and $DNAJB8_{F148S\_F151S}$ (teal) at two time points: 0 h (solid lines) and 30 h (dashed lines).

$DNAJB8_{WT}$ and $DNAJB8_{S3}$ are the same data as shown in Figure 1C (grey box). The size distribution of F148S matched the size of $DNAJB8_{WT}$ at t=0 h, with $DNAJB8_{F148S}$ showing evidence of self-assembly after t=30h. $DNAJB8_{F151S}$ and $DNAJB8_{F148S\_F151S}$ both remained monomeric throughout the experiment with average $R_h$ values indicating sufficient burial of hydrophobic surfaces. (**C**) Western blot of lysates from HEK293T cells expressing $DNAJB8_{WT}$-mRuby3, $DNAJB8_{S3}$-mRuby3, $DNAJB8_{F148S}$-mRuby3, $DNAJB8_{F151S}$-mRuby3, and $DNAJB8_{F148S\_F151S}$-mRuby3 run on native PAGE and treated with α-DNAJB8. Regions corresponding to monomers, low molecular weight oligomers, and high molecular weight oligomers are marked. (**D**) Transient expression of mRuby3 (red), $DNAJB8_{WT}$-mRuby3 (blue), $DNAJB8_{S3}$-mRuby3 (cyan), $DNAJB8_{F148S}$-mRuby3 (lime), $DNAJB8_{F151S}$-mRuby3 (purple) and $DNAJB8_{F148S\_F151S}$-mRuby3 (teal) in HEK293T cells. Data shown for mRuby3, DNAJB8, and DNAJB8 S3 was previously shown in Figure S1A (gray box). Expression is quantified by averaging the signal intensities across all captured images and normalized per the number of cells. These data show that changes in number of puncta observed are independent of expression. (**E**) Data shown for mRuby3, $DNAJB8_{WT}$, and $DNAJB8_{S3}$ was previously shown in Figure S1C (gray box) CellTiter-Glo luminescence for cell lines listed above. These data show that changes in puncta are independent of cell viability. (**F**) Data shown for mRuby3, $DNAJB8_{WT}$, and $DNAJB8_{S3}$ was previously shown in Figure S1C (gray box). Average diameter of puncta counted by CellProfiler across different cell lines. All cell line proteins are the same as in panel **A**. On average, most puncta detected in cells were around 1µm in diameter, regardless of mutations to DNAJB8. (**G**) Image of cells expressing $DNAJB8_{F148S}$-mRuby3 (red) with nuclei stained with DAPI (blue). Expression of mRuby3-tagged $DNAJB8_{F148S}$ is ubiquitous with protein found in both the cytoplasm and nucleus. However, inclusions are still observed for this mutant. Scale bar= 10µm. (**H**) Image of cells expressing $DNAJB8_{F151S}$-mRuby3 (red) with nuclei stained with DAPI (blue). Expression of mRuby3-tagged $DNAJB8_{F151S}$ is ubiquitous with protein found in both the cytoplasm and nucleus. Inclusions for this mutant are rare across the cell populations. Scale bar= 10µm. (**I**) Image of cells expressing $DNAJB8_{F148S\_F151S}$-mRuby3 (red) with nuclei stained with DAPI (blue). Expression of mRuby3-tagged

DNAJB8$_{F148S_151S}$ is ubiquitous with protein found in both the cytoplasm and nucleus. Inclusions for this mutant are rare across the cell populations. Scale bar= 10µm.

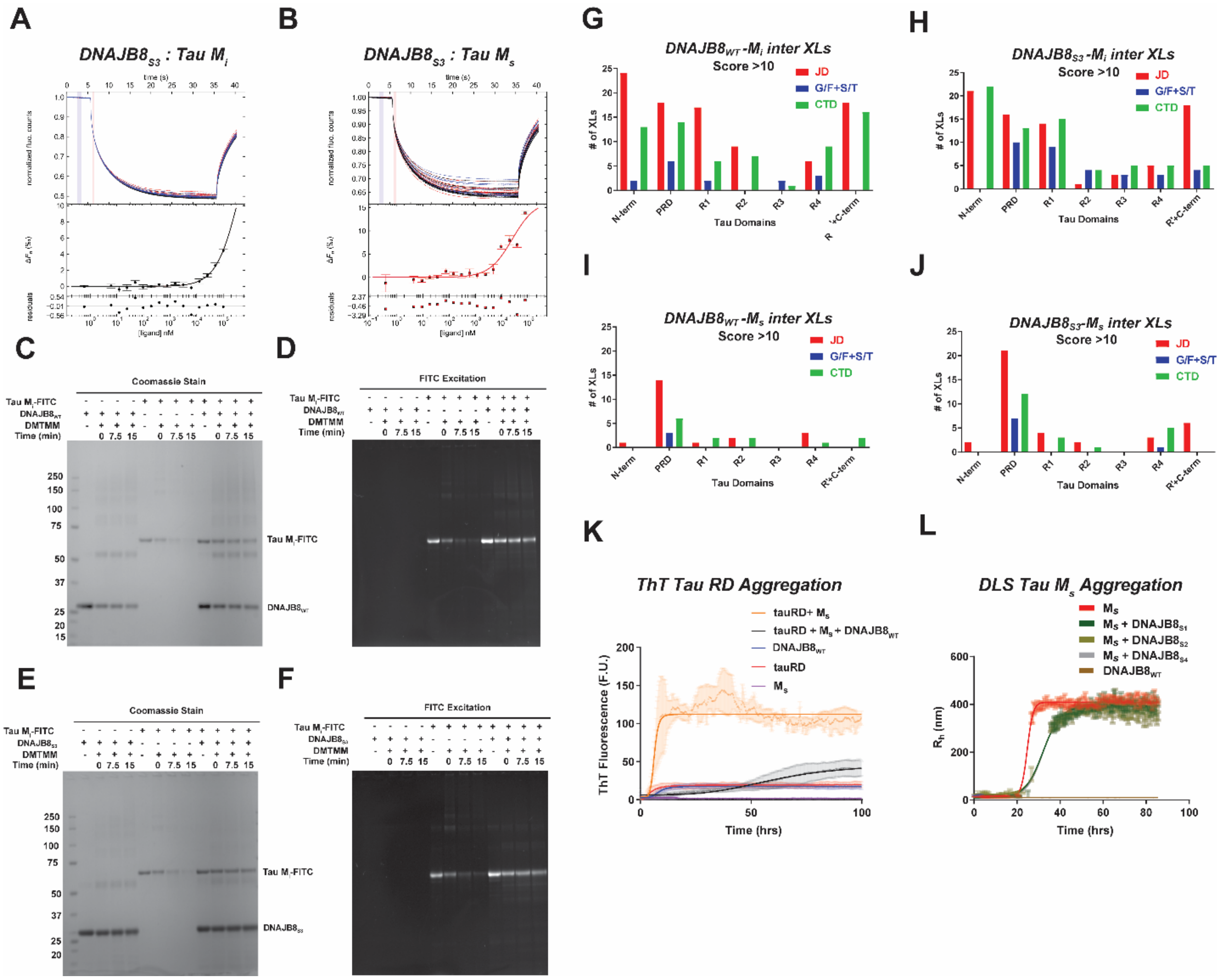


**Figure S4.** ***In vitro* activity of DNAJB8 against tau aggregation.** (**A**) Microscale thermophoresis where DNAJB8$_{S3}$ is titrated on a 2-fold step-wise scale to a fixed concentration of 1µM tau $M_i$. The curve was fitted to a 1:1 stoichiometry using PALMIST and $K_d$ was calculated to be >400µM. (**B**) Microscale thermophoresis where DNAJB8$_{S3}$ is titrated on a 2-fold step-wise scale to a fixed concentration of 1µM tau $M_s$. The curve was fitted to a 1:1 stoichiometry using PALMIST and $K_d$ was calculated to be >400µM. (**C**) SDS-PAGE imaged stained with Coomassie of protein samples crosslinked with DMTMM and quenched after 0 min, 7.5 min, or 15 min. At 20µM, DNAJB8$_{WT}$ does not show rapid self-assembly as opposed to higher concentrations. $M_i$ rapidly begins to aggregate upon addition of crosslinker with aggregates visible at the top of the gel. When $M_i$ is preincubated with DNAJB8$_{WT}$, there is a greater recovery of $M_i$ monomer even after 30 min of exposure to DMTMM. (**D**) The same SDS-PAGE as in panel **c**, but imaged for FITC fluorescence before the addition of coomassie stain. Here, only the labelled tau $M_i$-FITC bands are visible and all bands containing tau $M_i$-FITC are visible. This confirms that the retained monomer in the DNAJB8$_{WT}$ with tau $M_i$ samples is tau $M_i$, while also showing that the new observed band at ~80-90kDa contains tau $M_i$-FITC as one of its components. (**E**) SDS-PAGE imaged stained with Coomassie of protein samples crosslinked with DMTMM and quenched after 0 min, 7.5 min, or 15 min. DNAJB8$_{S3}$ shows the same effect on the gel as DNAJB8$_{WT}$, where when mixed with tau $M_i$ before adding DMTMM, there is a greater retention of tau $M_i$ monomer over time as opposed to tau $M_i$ by itself. (**F**) The same SDS-PAGE as in panel **E**, but imaged for FITC fluorescence before the addition of coomassie stain. The same effect on tau monomer retention is observed for DNAJB8$_{S3}$ as was observed for DNAJB8$_{WT}$. In addition, the same new tau-containing band at ~80-90kDa is observed when tau $M_i$ is in the presence of DNAJB8$_{S3}$ and DMTMM. (**G**) Histogram of total low-resolution (ID Score >10) intermolecular DMTMM crosslinks between DNAJB8$_{WT}$ domains (JD=red, G/F and S/T=blue, and CTD=green) and tau $M_i$ domains (x-axis). The distribution of cross-link pairs shows the JD and CTD are primarily interacting with tau, but in a heterogenous fashion across the protein. (**H**) Histogram of total low-resolution intermolecular DMTMM crosslinks between DNAJB8$_{S3}$ domains and tau $M_i$ domains. The distribution of cross-link pairs shows the JD and CTD are primarily interacting with tau, but in a heterogenous fashion across the protein. A few more contacts are observed with the disordered G/F and S/T-rich domains, but otherwise DNAJB8$_{S3}$ is interacting with tau $M_i$ in a similar manner to DNAJB8$_{WT}$. (**I**) Histogram of total low-resolution intermolecular DMTMM crosslinks between DNAJB8$_{WT}$ domains and tau $M_s$ domains. These data show that the JD and CTD of DNAJB8$_{WT}$ continue to primarily participate in crosslinks to tau. However, there is a clear spike in crosslinks localized to the tau PRD. (**J**) Histogram of total low-resolution intermolecular DMTMM crosslinks between DNAJB8$_{S3}$ domains and tau $M_s$ domains. Similar to the data between DNAJB8$_{WT}$:tau $M_s$, DNAJB8$_{S3}$:tau $M_s$ shows a distinct majority of contacts between the JD/CTD of DNAJB8$_{S3}$ and the tau $M_s$ PRD. (**K**) tau $M_s$ induced aggregation assay of tauRD using ThT fluorescence. Control curves are shown for 10µM DNAJB8$_{WT}$ (blue), 5µM tauRD (red), and 0.05µM tau $M_s$ to show that each individual protein does not self-assemble at the given concentrations over the 100h time-course. Tau $M_s$ seeded tauRD (orange) showed rapid tau aggregation after 5h ($t_{1/2max}$= 6.3 ± 0.2 hours), while tau $M_s$ seeded tauRD that was preincubated with $DNAJB8_{WT}$ (black) showed a long, delayed increase in ThT signal over time ($t_{1/2max}$= 61.6 ± 2.9 hours). (**L**) Dynamic light scattering time course experiment reporting average $R_h$ as time points for $DNAJB8_{WT}$ (brown), tau $M_s$ (red), $DNAJB8_{S1}$ and tau $M_s$ (green), $DNAJB8_{S2}$ and tau $M_s$ (gold), and $DNAJB8_{S4}$ and tau $M_s$ (gray). Each DNAJB8 mutant was mixed with tau $M_s$ at a 100:1 ratio. At these conditions, all of these mutants are able to slow the rate of tau aggregation by ~20 hrs, matching the $DNAJB8_{WT}$ results.

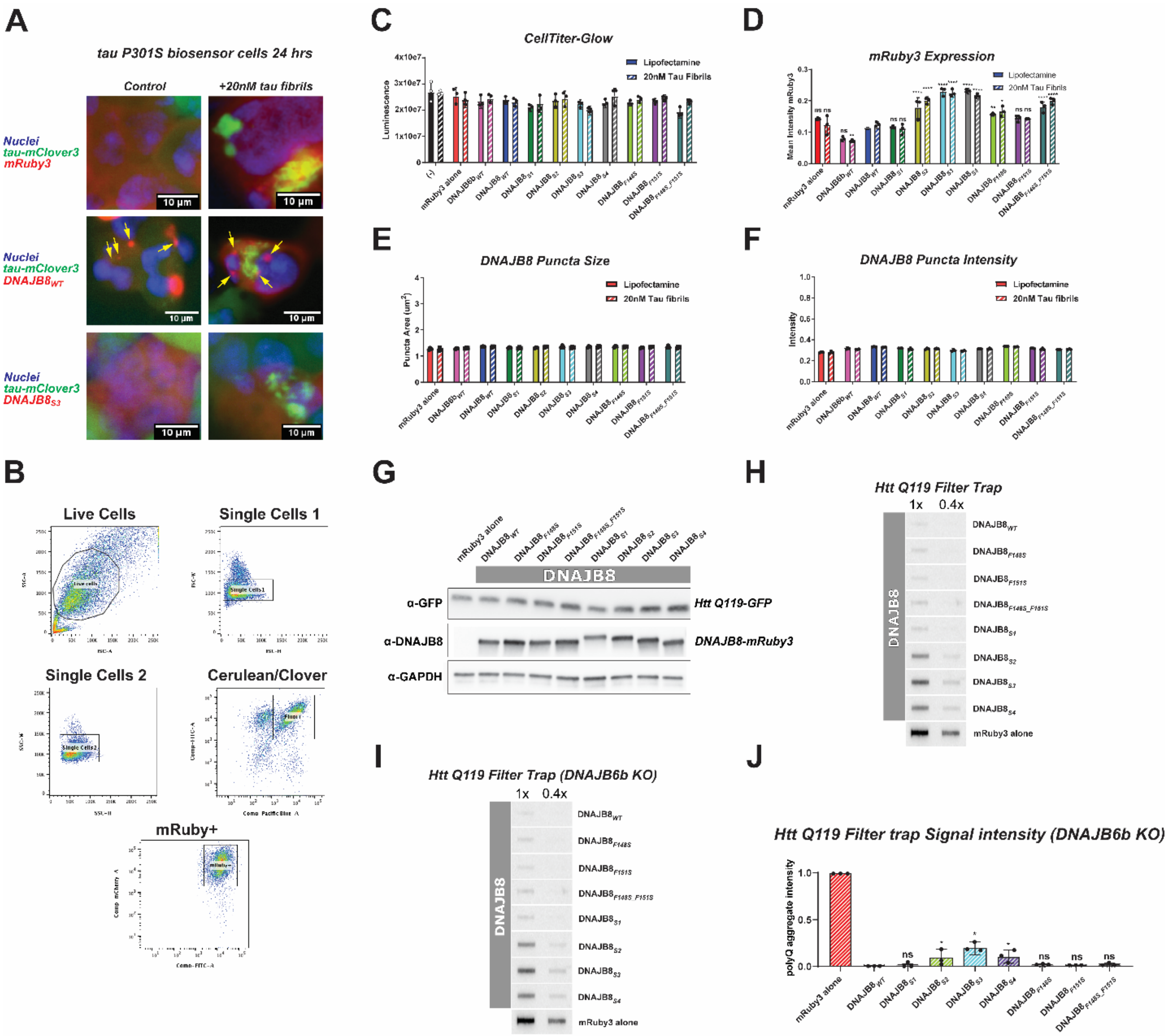

**Figure S5. Tau and polyQ anti-aggregation activity of WT and mutant DNAJB8 in cells.** (**A**) Images of HEK293T tau biosensor cells showing expression of tauRD-mClover3 (green), mRuby3 (red), DNAJB8$_{WT}$-mRuby3 (red), DNAJB8$_{S3}$-mRuby3 (red), and nuclei stained with DRAQ5 (blue). In the unseeded control lines, tauRD-mClover3, mRuby3, and DNAJB8$_{S3}$-mRuby3 signals overlap throughout the entire cell, with DNAJB8$_{WT}$-mRuby3 only overlapping in the cytoplasm. DNAJB8$_{WT}$-mRuby3 puncta are visible in both control and seeded conditions (yellow arrows). When seeded, tauRD-mClover3 accumulates as large aggregates in the cells, but does not co-localize with DNAJB8$_{WT}$-mRuby3 puncta, nor co-localize with ubiquitously expressed DNAJB8$_{S3}$-mRuby3. (**B**) Gates were drawn with standard flow cytometry methodology for Live cell population, followed by FSC single cells and SSC single cells. Single cells expressing tau P301S RD mCerulean/mClover were isolated, a sub-gate was used to select cells DNAJB8 tagged to mRuby (mRuby+). FRETClover/Cerulean was calculated from this population as was shown previously[56, 78] (**C**) CellTiter-Glo luminescence assay of tau biosensor lines that were treated with lipofectamine or 20nM tau fibrils 24 hours after initial transfection of DNAJB8-mRuby3 containing plasmids. Under these conditions, the uptake of $DNAJB8_{WT}$ or its mutants had no impact on the viability of the cell populations, nor did the 20nM tau fibril seeding significantly impact cell viability. These data show that the results in Figure 5 are not influenced by significant cell death. (**D**) mRuby3 signal quantified across all cell populations based on values generated by image analysis in CellProfiler v.4.2.1. Significance calculated using two-way ANOVA relative to $DNAJB8_{WT}$ unseeded and seeded lines. * $p<0.1$, ** $p<0.01$, *** $p<0.001$, **** $p<0.0001$ (**E**) Calculated average mRuby3 puncta size per cell population. Average puncta area remains consistent across all DNAJB8 samples. (**F**) Calculated average mRuby3 puncta intensity per cell population. (**G**) Western blot of DNAJB8-mRuby3 expression in HEK293T HttEx1 Q119 cell lines. Samples were treated with α-GFP for Htt Q119-GFP, α-DNAJB8 to blot for DNAJB8-mRuby3, or α-GAPDH for loading control. (**H**) Raw FTA blot of HttEx1 Q119-GFP in cells transfected with DNAJB8-mRuby3 mutants. Band intensities across 1x and 0.4x dilution were quantified and plotted in Figure 5g to characterize DNAJB8 chaperone activity. (**I**) Raw FTA blot of HttEx1 Q119-GFP in DNAJB6b KO cells transfected with DNAJB8-mRuby3 mutants. Endogenous DNAJB6b expression was shut off in cells before the experiment to remove any anti-polyQ aggregation effects driven by DNAJB6b. Observed HttEx1 Q119 bands are a reflection of only DNAJB8 activity in cells. (**J**) FTA results showing HttEx1 Q119 aggregation for DNAJB6b KO cells expressing mRuby3 (red), $DNAJB8_{WT}$-mRuby3 (blue), $DNAJB8_{S1}$-mRuby3 (green), $DNAJB8_{S2}$-mRuby3 (yellow), $DNAJB8_{S3}$-mRuby3 (cyan), $DNAJB8_{S4}$-mRuby3 (grey), $DNAJB8_{F148S}$-mRuby3 (lime), $DNAJB8_{F151S}$-mRuby3 (purple), and $DNAJB8_{F148S\_F151S}$-mRuby3 (teal). All values were normalized to the mRuby3 negative control. A small loss of DNAJB8 activity was reported for $DNAJB8_{S2}$-mRuby3,

$DNAJB8_{S3}$-mRuby3, and $DNAJB8_{S4}$-mRuby3 expressing cells. Statistics were calculated using Welch's T test for each pair relative to $DNAJB8_{WT}$, where * $p<0.1$.
